# Controlling False Positive Rates in Methods for Differential Gene Expression Analysis using RNA-Seq Data

**DOI:** 10.1101/018739

**Authors:** David M. Rocke, Luyao Ruan, J. Jared Gossett, Blythe Durbin-Johnson, Sharon Aviran

## Abstract

We review existing methods for the analysis of RNA-Seq data and place them in a common framework of a sequence of tasks that are usually part of the process. We show that many existing methods produce large numbers of false positives in cases where the null hypothesis is true by construction and where actual data from RNA-Seq studies are used, as opposed to simulations that make specific assumptions about the nature of the data. We show that some of those mathematical assumptions about the data likely are one of the causes of the false positives, and define a general structure that is not apparently subject to these problems. The best performance was shown by limma-voom and by some simple methods composed of easily understandable steps.

## 1 Introduction and Background

### 1.1 Gene Expression

Transcriptome profiling is a powerful means of characterizing a cell’s state and as such, is key to elucidating cellular processes, dissecting genotype-phenotype relationships, and understanding health and disease. As genes are expressed at levels that can vary greatly both within a cellular sample and between samples, it is imperative to not only quantify these expression levels robustly but to also detect and assess between-sample differences with high fidelity.

For nearly two decades, cDNA microarrays have been the platform of choice for profiling complex mixtures of RNAs sampled from the transcriptome (Bertone, Stolc [1], Subramanian, Tamayo [2]). Microarray technology transformed and accelerated gene expression studies by enabling large-scale and genome-wide measurements via parallel, automated, and cost-efficient hybridization-based detection of nucleic acids. While it accurately quantifies relative prevalence of transcripts that derive from a set of pre-identified genes, this technology also suffers major limitations. Primarily, it cannot support exploratory studies or facilitate *de novo* discoveries, as it requires *a priori* design and fabrication of the array with the genes that are to be probed. Such considerations essentially preclude unannotated genes or unknown splice variants from being detected as well as limit the number of variants that a single experiment can assay. Other issues include limited dynamic range and imperfections in signal acquisition, which warrant dedicated and non-trivial analysis methodology.

A fundamentally new approach to large-scale and comprehensive RNA analysis has emerged with the recent advent of next-generation DNA sequencers (Wold and Myers [3]). While this technological advance was initially directed at massive genome sequencing, it was soon harnessed to provide sequencing-based RNA transcript quantification, dubbed RNA-Seq (Mortazavi, Williams [4], Nagalakshmi, Wang [5]). RNA-Seq was rapidly adopted and standardized due to the assay’s simplicity, affordability, enhanced information content, and scalability; it is standard practice in molecular biology and biomedical research (Van Keuren-Jensen, Keats [6] Li, Tighe [7]). This paradigm shift clearly contributed to the scaling trajectory of gene expression studies, but most importantly, it expanded their scope by allowing detection of transcripts *de novo* and at very low abundances. Yet, such unprecedented level of detail also presents multiple new informatics challenges, in particular with respect to transcript reconstruction, quantification, and comparative or differential analysis. In this review, we focus on the latter challenge—an area still early in its intellectual development and which is changing rapidly. We highlight state-of-the-art methods for whole-genome differential expression analysis from RNA-Seq data by placing them within a common framework that allows comparison of their performance. We also review a number of recent studies comparing these methods in terms of false positives and sensitivity, and add additional results of our own.

### 1.2 Previous Comparisons of Packages for Differential Expression Analysis

A number of recent papers have compared the power and type I error rates of popular packages for analysis of RNA-Seq Data, with varying conclusions. Soneson and Delorenzi [8], Kvam, Liu [9], and van de Wiel, Neerincx [10] compared the null performance of several methods and found poor control of false discovery rate (FDR) in general. Robles, Qureshi [11] likewise compared 3 methods and found inflated type I error rates for some methods and conservative performance from others. Reeb and Steibel [12] compared 3 methods using “plasmodes” (resampled data) and found inflated type I error rates for small significance levels. Guo, Li [13] compare 6 methods and conclude that all “suffer from over-sensitivity”. However, Rapaport, Khanin [14] compared 7 different methods and concluded that most methods provide good FDR control at the 5% level. Finally, Nookaew, Papini [15] compared 5 different methods with each other and with microarray data and concluded that agreement was good between methods and platforms, and Guo, Li [13] concluded that agreement between the methods they compared was “reasonable”.

Our comparison is limited to five commonly used methods implemented in Bioconductor (Gentleman, Carey [16]): edgeR (Robinson, McCarthy [17], McCarthy, Chen [18], Robinson and Smyth [19], Robinson and Smyth [20]), edgeR-robust (Zhou, Lindsay [21]), limma-voom (Smyth [22], Law, Chen [23], Ritchie, Phipson [24]), DESeq (Anders and Huber [25]), and DESeq2 (Love, Huber [26]). Many other methods exist, including but not limited to: Cuffdiff (Trapnell, Williams [27]), Cuffdiff2 (Trapnell, Hendrickson [28]), NBPSeq (Di, Schafer [29]), TSPM (Auer and Doerge [30]), baySeq (Hardcastle and Kelly [31]), EBSeq (Leng, Dawson [32]), NOISeq (Tarazona, García-Alcalde [33]), SAMseq (Li and Tibshirani [34]), ShrinkSeq (Van De Wiel, Leday [35]), DEGSeq (Wang, Feng [36]), BBSeq (Zhou, Xia [37]), FDM (Singh, Orellana [38]), RSEM (Li and Dewey [39]), Myrna (Langmead, Hansen [40]), PANDORA (Moulos and Hatzis [41]), ALDEx2 (Fernandes, Reid [42]), PoissonSeq (Li, Witten [43]), and GPSeq (Srivastava and Chen [44]). The methods we chose to examine in detail seemed to be the most commonly used and are easy to run in R. We also ran the packages with the default options, since that is likely what most users do, and since otherwise the combinatorial complexity can get out of hand. We provide code that can be adapted to any method that runs in R and applied to the publicly available data sets we used, as well as others.

### 1.3 Overview of Methods for Differential Expression Analysis with RNA-Seq Data

Although the array of seemingly complex methods and packages suggests that this is a complicated and difficult task, at its core differential expression analysis is simple. If one knows how to test treatment vs. control when a single measurement has been made per sample, then most of what is needed for DE analysis has been accomplished. Most of the task consists of testing each gene (as a row in the data matrix) and returning a p-value as a measure of statistical significance (or perform other statistical tasks). There are a few other components that are or may be needed: filtering, normalization, improved variance estimation, and adjustment for multiple comparisons, but a method of testing each gene is all that is really required for a feasible method for determining the statistical significance of a set of genes in DE analysis for RNA-Seq. In fact, good methods for analysis of gene expression arrays, proteomics by mass spectrometry, and metabolomics by mass spectrometry share almost all the same features. In this section, we describe these components and some alternatives. We do not describe in detail the choices and methods and all the options in the packages commonly used for RNA-Seq, though we will outline the general approach in each one that we examine.

#### Statistical Tests

Perhaps the most fundamental choice is the basic statistical test used for each gene. Many packages and methods, including DESeq (Anders and Huber [25], [45]), DESeq2 (Love, Huber [26]), edgeR (Robinson, McCarthy [17], McCarthy, Chen [18], Robinson and Smyth [19], Robinson and Smyth [20]) and edgeR-robust (Zhou, Lindsay [21]) are based on a negative binomial model for the counts in the expression matrix. The mathematics of this is detailed in most of the papers supporting these methods, but the rationale seems to be something like this:

1. RNA-Seq data are in the form of counts.
2. The simplest model for count data is the Poisson, in which the mean and the variance are the same.
3. But in most cases, the data are over-dispersed, meaning that the variance is greater than the mean. This may occur when individual observations for a given gene under given conditions are each Poisson, but with a mean that varies from observation to observation.
4. Perhaps the simplest distribution for over-dispersed count data is the negative binomial, which is obtained when each observation is Poisson but the mean varies from sample to sample following a gamma distribution.

One consequence of this assumption is a relationship between the mean and the variance. If a random variable X has a negative binomial distribution with mean μ and variance σ^2^, then σ^2^ = μ + θμ^2^, where θ can be called the dispersion parameter. So, in a comparison of treatment vs. control, we need to estimate two means, μ_1_ and μ_2_, and the dispersion parameter θ, which then determines the two variances 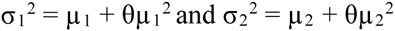. It is also commonly assumed that the parameter θ is common to all genes, at least approximately. In general, then, tests that are based on the negative binomial distribution involve an assumption that there is a mean-variance relationship that is shared by all the measurements of a particular gene from all the samples. But over-dispersion does not itself imply that the distribution is negative binomial, only that it is not Poisson, so these assumptions may not be correct in practice.

The limma package (Smyth [22]), originally developed for gene expression microarray analysis, is based on fitting a standard linear model for each gene. By default, in a standard linear model, the error variance is the same for all observations from the same gene, for treatment or control. After a log-like transformation (Durbin, Hardin [46]; Durbin and Rocke [47]; Durbin and Rocke [48]; Rocke and Durbin [49]; Rocke and Durbin [50]), this might be a plausible assumption for microarray data, though care in the choice of the transformation is important. For RNA-Seq data also, the variance tends to rise with the mean in a very marked fashion, so that the assumption of equal variance cannot hold in general without a transformation. Taking logarithms can help with this problem, but it fails for low counts, especially zero counts. For the basic treatment vs. control comparison, limma conducts what is actually a pooled two-sample t-test, which assumes equal variance. In order to handle the problem of variances trending with the mean, the voom function (Law, Chen [23]) was written as a front end for limma, yielding a method called limma-voom. The voom function calculates variance-based weights that, together with the log transformation in limma, are meant to stabilize the variance.

In addition, we consider two simple methods that can be applied to the data from a single gene. Neither of these is specifically part of a package, but both arise from standard statistical methods. Neither of them uses the variance shrinkage methods described below, though the simple methods could also use variance shrinkage if desired. The first of these is the two-sample t-test that does not assume equal variance (Welch [51]). The second test arises from the generalized linear model regression formulation with the negative binomial distribution as implemented in the R function glm.nb() in the MASS package (Ripley [52]). This is the likelihood ratio test (Neyman and Pearson [53]) which compares the model to a simpler model, which is similar to the ANOVA F-test in ordinary linear regression (Snedecor [54]). In the case of a two-condition experiment, we compare a model with the condition as a predictive factor to one that has only the mean as a predictor.

We will also consider the application of a transformation to the counts before applying a test. As described in Bartlett’s 1947 paper (Bartlett [55]), once a variance function (relationship between the variance and the mean) has been defined, one can derive a transformation that approximately stabilizes the variance. This is to some extent described in the vignette for the DESeq package (Anders and Huber [25]). After some derivation it can be shown that if σ^2^ = μ + θμ^2^, then the transformation

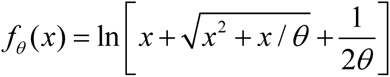

approximately stabilizes the variance, meaning that the variance no longer depends systematically on the mean. Thus, after transformation, we can use methods that don’t explicitly model the variance as a function of the mean, such as linear regression, the analysis of variance, or in the simple two-condition case, the pooled t-test. Figures 1–5 show the mean variance relationship for the Montgomery/Pickrell data set, which is described later, containing RNA Seq data from 60 individuals of European heritage and 69 of African heritage ([56], [57]). The plots show the mean-variance relationship with (respectively) no transformation, the log transformation, and three values of θ, one too small, one too large, and one that seems about right. One way to estimate the best value of θ is to regress the gene-and-condition-specific variance on the gene-and-condition-specific mean, and find a value of θ for which the variance neither increases systematically with the mean nor decreases systematically. We can operationalize this by finding a value of θ for which the slope is zero, and this is how the “about right” value was estimated.

**Figure 1.**
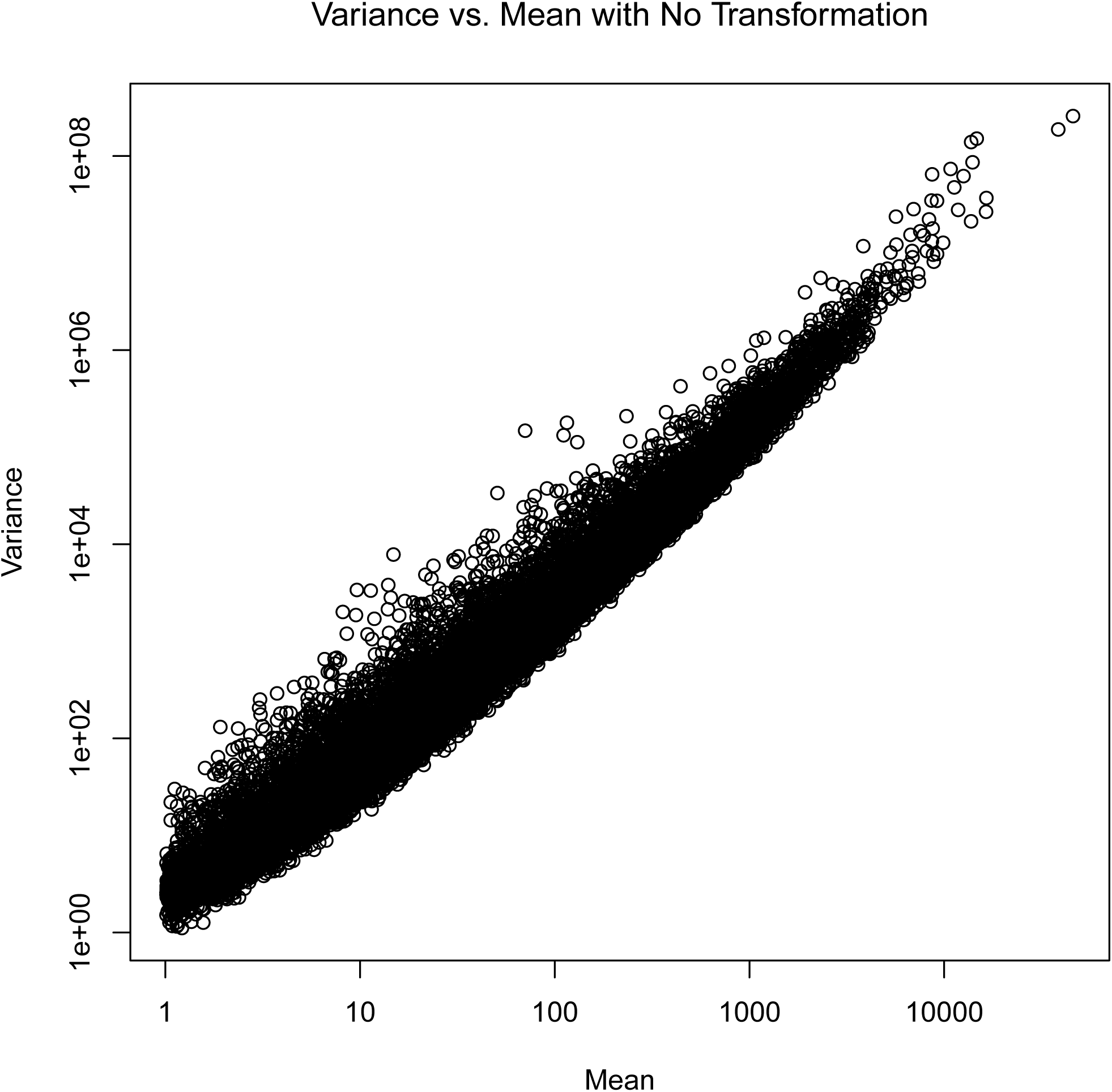
Group-and gene-specific variances plotted against the means without transformation (axes on the log scale).

**Figure 2.**
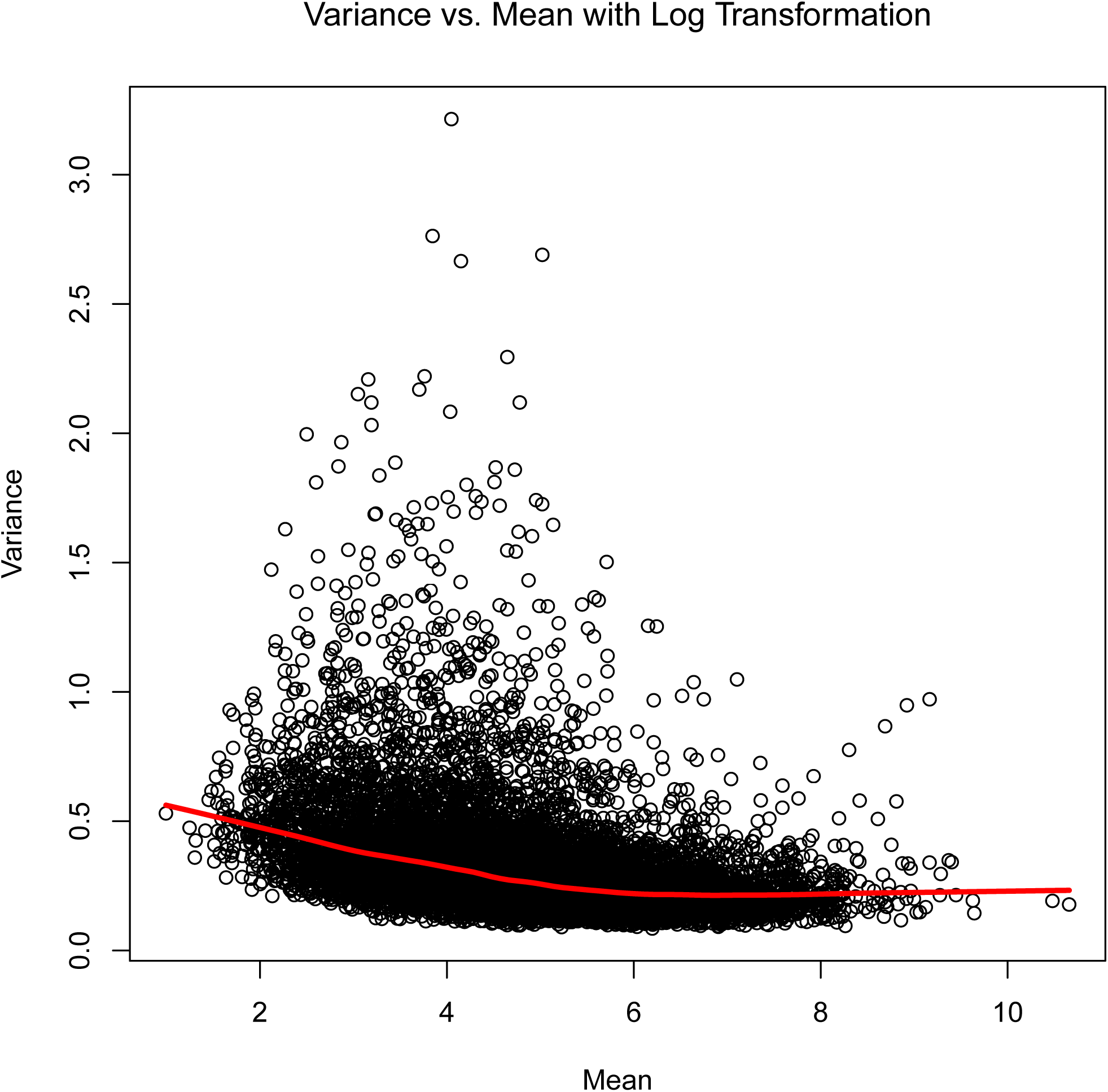
Group- and gene-specific variances plotted against the means after a log transformation.

**Figure 3.**
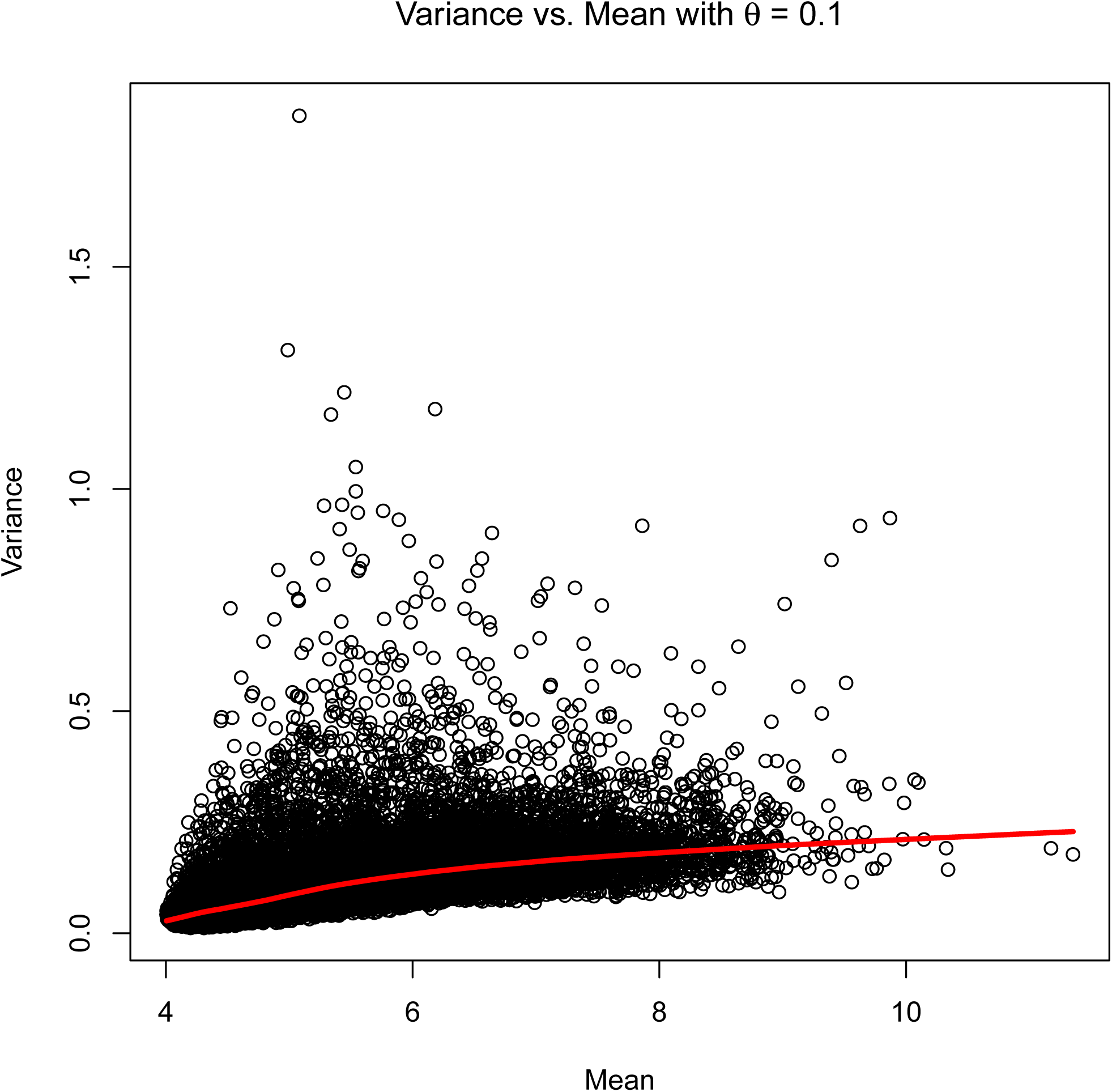
Group- and gene-specific variances plotted against the means after a variance-stabilizing transformation with θ = 0.1.

**Figure 4.**
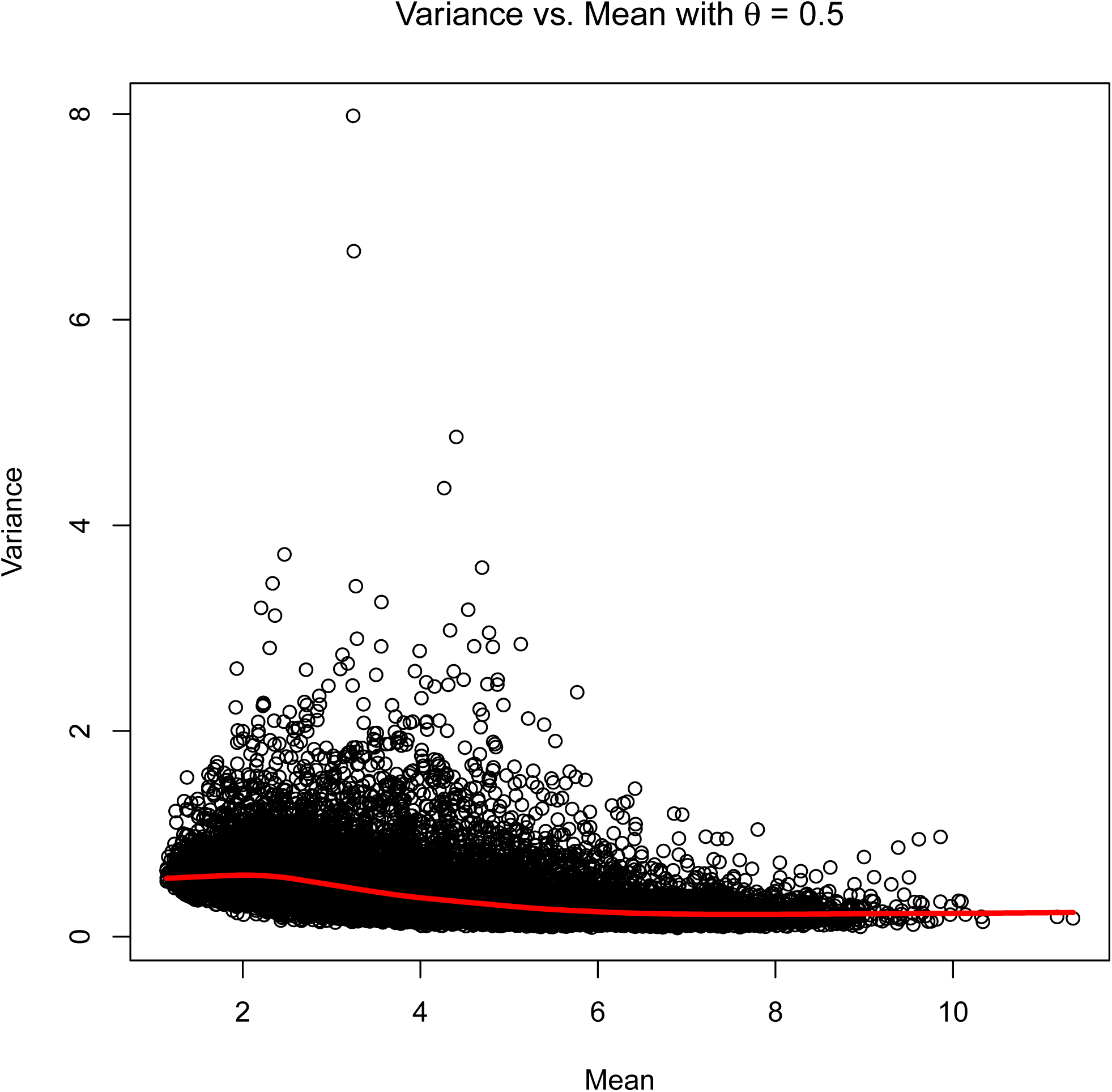
Group- and gene-specific variances plotted against the means after a variance-stabilizing transformation with θ = 0.5.

**Figure 5.**
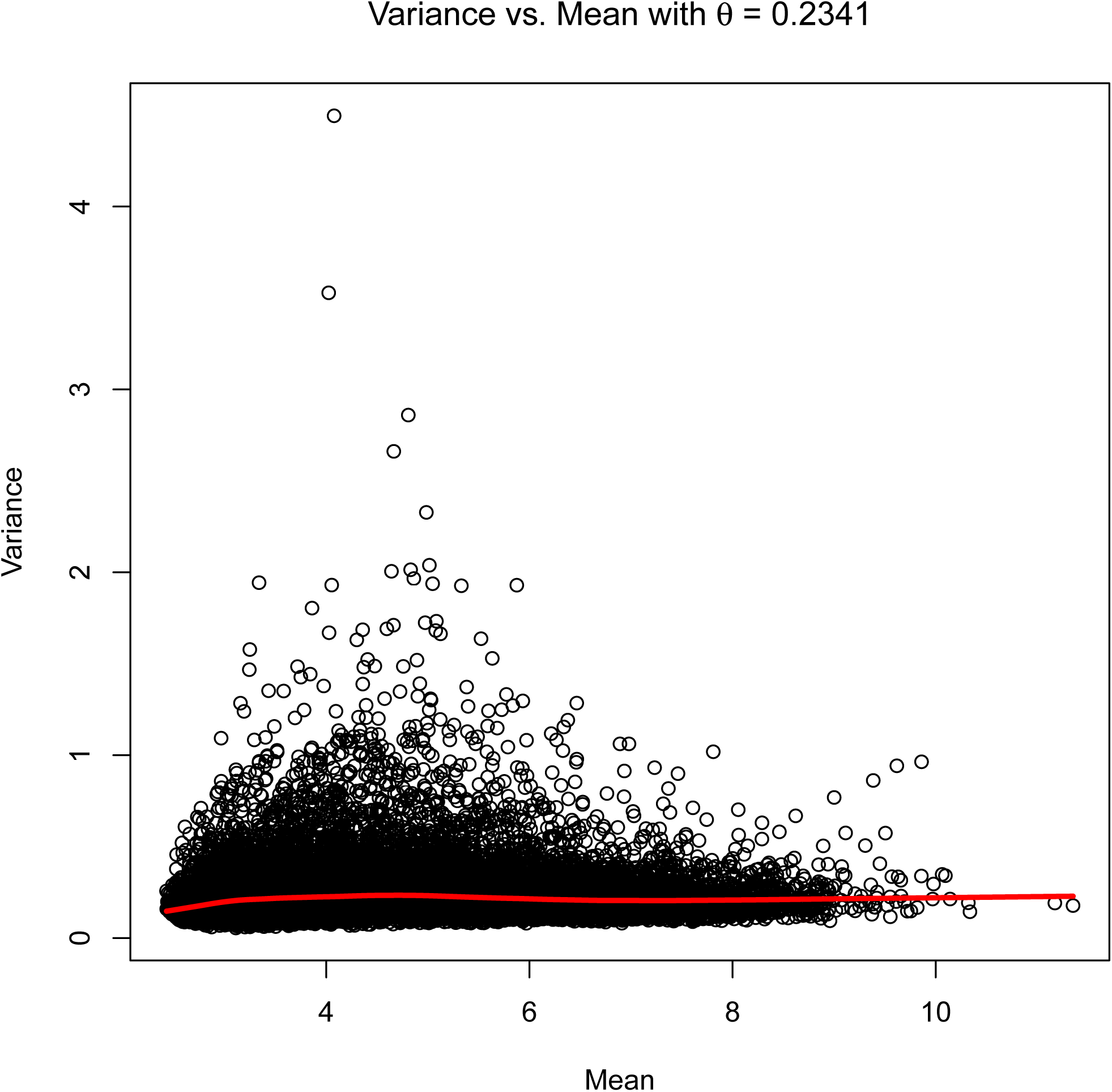
Group- and gene-specific variances plotted against the means after a variance-stabilizing transformation with θ = 0.2341.

#### Filtering

By filtering, we mean removing from consideration genes that do not satisfy some criterion for large enough counts. If we have, for example, 10 samples in condition 1 and 11 in condition 2, we might require that the sum of the counts across all 21 samples is larger than some lower limit such as 10 or 21, or that the number of samples with zero or very low counts is smaller than some limit such as 10. On the one hand, this may be needed if the method performs badly when the counts are low. On the other hand, this means that detection of differential expression is not possible for these genes, and of course some genes that are very important biologically may not have transcripts that are present in large quantities.

#### Normalization

Normalization means adjusting all of the values from a given sample by a quantity or quantities that are common to all the genes in the sample. In RNA-Seq, this is often referred to as library size, meaning some total fragment count, but similar procedures are also used in other high-throughput gene expression methods, such as gene expression arrays. It is commonly assumed that normalization is always required, but in fact this is an empirical issue. If the count in a two-group experiment is assumed to follow

*k*_*ijk*_ = *μ*_*i*_ + *β*_*ij*_ + *γ*_*jk*_ + ϵ_*ijk*_ from gene *i* in condition *j* and biological replicate *k*

*μ*_*i*_ = average expression of gene *i* across all conditions.

*β*_*ij*_ = effect of condition *j* on the expression of gene *i*

*γ*_*jk*_ = effect of sample *jk* on measured expression of all genes

ϵ_*ijk*_ = measurement error

then the sample effect γ increases the variance of the error, and induces a correlation across genes. If we try to remove this with an estimate *g* of γ, then we might possibly reduce the error variance, but depending on the bias and variance of *g*, we might also increase it. Nonetheless, normalization seems virtually always to be used. Each package has one or more such methods—often geometric mean normalization is chosen—but for the simple estimators we use median normalization in which each value in a sample is divided by the median count in that sample and then multiplied by the median of all the sample medians.

#### Variance Estimation

With large sample sizes, we hardly need anything else to perform differential analysis. But often the sample sizes are small, leading to low power. This lead to the development of a method using empirical Bayes to “borrow strength” across genes (Wright and Simon [58], Smyth [59], Baldi and Long [60]). This is most simply explained in a context in which the variance does not depend systematically on the mean.

Suppose we have a linear model for each gene. This could be a two-condition comparison (such as treatment vs. control) or a more complex model such as linear regression or multi-factor analysis of variance. Hypothesis tests for these models all depend on an estimate of the residual mean square error. For example, if we model the response as having one mean under the treatment group and one mean under the control group, then the t-test, as with almost any other statistical test, compares the difference of the means to the variability within the groups (in this case the pooled standard deviation). Given the sample means and standard deviations of treatment and control, and assuming for simplicity that there are n samples in each group,

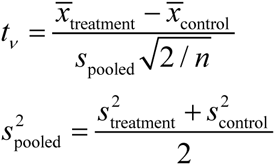

If n is small (for example, n = 3), then the estimate of the variance is poor, with (in this case) only 4 degrees of freedom (df). If we are looking for significance at a p-value of, say, 10^-4^, then with a large number of df, a test statistic of 3.89 is sufficient, but with 4 df, the test statistic would need to be 15.54, a much higher hurdle to surmount.

In the empirical Bayes method, we try to improve these variance estimates by higher level modeling. If we have 10,000 genes and 10,000 statistical tests, we have 10,000 residual variances to estimate. This is done somewhat differently in methods that are not based on linear models, but the principles are similar. For each gene we have an observed residual mean square error 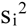, each with degrees of freedom ν. Then (under normality), each of these is a multiple of a chi-squared random variable:

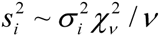

Each gene has a true error variance 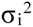 and an estimate of it 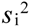. If ν = 4, then the critical value for p = 10^-4^ is 15.54. If we knew σ, then the critical value is only 3.71. If we could somehow improve the estimate of σ so that it behaved like (say) an 8-df estimate, then the critical level would be 7.12, a huge improvement.

The method of improving the estimates rests on thinking of the true error variances of the linear models as random variables from a (prior) distribution. For technical reasons, we use an inverse gamma distribution for the prior, meaning that the distribution of the reciprocal of the variance is a gamma distribution, which is the same as a chi-squared. This distribution has two parameters, for example the reciprocal mathematical expectation 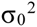 and the degrees of freedom ν_0_. It turns out then that the posterior distribution of the true error variance for a specific gene is also inverse gamma with posterior mean

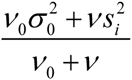

which is a weighted average of the prior variance and the observed variance from gene *i*, each weighted by the degrees of freedom. The parameters of the prior can be estimated as described in Wright and Simon [58] or Smyth [59]. This approach is called *empirical* Bayes, because the prior distribution does not come from opinion, or from a representation of prior lack of information, but is estimated from the data. We can also describe this as “shrinking” the individual sample variances towards a kind of average of the sample variances (technically the prior variance is approximately the reciprocal of the average of the reciprocal individual error variances). This is essentially the procedure used in limma.

When the variance trends with the mean, we can then shrink the variances towards the expected variance given the mean, or we can shrink the dispersion factor θ toward its mean. This is essentially the method used in DESeq2 and both edgeR variants. The package DESeq by default estimates the variance as the larger of the expected variance/dispersion and the observed variance/dispersion. This necessarily biases the variance up, causing a loss of statistical power.

### 1.4 Results of Application of Several Commonly Used Packages to Three Example Datasets

We first compare five commonly used packages on three data sets, the Bottomly data (Bottomly, Walter [61]), the Cheung data (Cheung, Nayak [62]), and the Pickrell/Montgomery (“MontPick”) data (Pickrell, Marioni [56], Montgomery, Sammeth [57]) as obtained from the recount site (http://bowtie-bio.sourceforge.net/recount/). The Bottomly data consist of RNA-Seq results from 21 samples of gene expression in the striatum in two strains of mice, 10 from the C7BL/6J strain and 11 from the DBA/2J strain. The Cheung data consist of RNA-Seq results from 41 human samples of B-cell gene expression from the HapMap project. These were from the CEPH (Utah residents with ancestry from northern and western Europe) (CEU) subsample and 17 of the samples were female and 24 were male. The MontPick data consist of RNA-Seq results from 129 human samples of B-cell gene expression from the HapMap project. There were 60 from the above-referenced CEU subset and 69 from the Yoruba people in Ibadan, Nigeria (YRI).

We applied each of the five packages/methods—DESeq, DESeq2, edgeR, edgeR-robust, and limma-voom—to each data set with the default options.1 We filtered the data to include only genes in which the mean of the counts across all the samples is at least 1. Table 1 shows the number of genes before and after filtering, the number of samples in each group, and the number of genes significant at a p-value cutoff of 10^-4^.

**Table 1.** Three data sets and the numbers of genes significant for differential expression for five methods with a p-value cutoff of 10^-4^.

| Data Set | Group 1 | Group 2 | Genes | Filtered Genes | Expected Under Null | DESeq | DESeq2 | edgeR | edgeR Robust | Limma Voom |
| --- | --- | --- | --- | --- | --- | --- | --- | --- | --- | --- |
| Bottomly | 10 C57BL/6J | 11 DBA/2J | 36536 | 11175 | 1.1175 | 327 | 573 | 529 | 639 | 385 |
| Cheung | 17 Female | 24 Male | 52580 | 9067 | 0.9067 | 5 | 11 | 21 | 25 | 7 |
| MontPick | 60 CEU | 69 YRI | 52580 | 8124 | 0.8124 | 1295 | 2762 | 2728 | 3150 | 2580 |

We can see that some methods produce many more apparently significant genes than others, with edgeR robust giving the largest number and with substantial differences between data sets, likely due in part to the number of truly differentially expressed genes. The differences are larger in the smaller data sets than in the larger ones as might be expected, because larger sample sizes can smooth out differences in the methodologies. In later sections of the paper, we try to estimate how many of these in each case are false positives and how many are reliable true positives, where by reliable we mean that the rate of “significant” genes in a data set with no true differences is not substantially greater than the nominal significance rate. In other words, we can’t just order the methods from best to worst according to the number of genes that the method says are differentially expressed, because then the best method would be to declare all genes to be differentially expressed. We must add the constraint that the fraction of genes declared significant, when there are truly no differences, must be approximately equal to the nominal significance level. At p = 0.0001, we should not declare more than about 0.01% of the genes significant if there are no true differences.

We chose the 10^-4^ level to make some allowance for multiple comparisons, but not to be so low as to be unduly restrictive. For comparison, with 10,000 genes, the cutoff with the Bonferroni method for a 5% test is 5×10^-6^. We did not use the FDR-adjusted p-values because each of those is a complex function of the entire vector of p-values, and this can make interpretation more difficult, but the raw p-values for a typical method in this data set for FDR-adjusted p-values of 5% are in the range of 10^-4^ and lower. Note that if the null hypothesis is always true, then all genes identified as significant are false positives. If we ask for a set of genes with FDR control at 5%, that means that the number of false positives should be no more than 5% of those identified as significant, and since all genes identified as significant are false positives, that means that an FDR-controlling method should never identify any genes as positive, at least on the average.

In every case, edgeR-robust has the largest number of significant genes and DESeq the fewest. The conservatism of DESeq in these cases is likely induced by the use of the maximum of the realized variance and the smoothed curve. Still, it is not clear which of these methods gives better results until we know how they perform under the null hypothesis of no difference.

## 2. Methods—Measuring Performance when There are no True Group Differences

When we analyze a data set, there may (depending on the package) be many steps in the analysis meant to normalize samples, borrow information from gene to gene, and so on, but the fundamental process is to compare for each gene the counts from the samples in one group to the counts from samples in the other (or in general a more complicated set of comparisons). Simply put, we want to see if the counts from one group are so much larger or smaller than the counts from the other group that the differences cannot easily be explained by variation within the group. We explicitly or implicitly treat the group 1 samples as more or less similar to each other and the group 2 samples as more or less similar to each other and then ask for each gene if the two groups differ from each other. This also implies that if we randomly selected say 3 of the samples from group 1 and 3 of the samples from group 2, then we would be making the same assumptions and asking the same questions. Of course, with smaller sample sizes, we would likely have less power to detect differential expression, but this would mirror the situation if we had not had the full number of samples to begin with but only 6, not an uncommon situation in RNA-Seq.

But what if we select 3 samples from group 1 at random and compare them with another randomly selected group of 3 samples from the same group? Since there are no systematic differences, we should find that the fraction significant is within chance range of the nominal significance level. With, say, 10,000 genes, we should have on the order of 1 gene significant at p = 10^-4^. Of course we could be unlucky and the 3 vs. 3 could have some factor that is more similar within these random groups than across them, but if we repeat the random selection a sufficient number of times, that should average out. Thus, if these methods are performing as advertised, when we conduct a “null” experiment like this, we should not have substantially more significant genes than is implied by the significance level.

We use this simple method to evaluate the null performance of the packages. For each of the three data sets, we conduct this experiment with 3, 4, or 5 in each subgroup of group 1 and then also for subgroups of group 2. For the Cheung data set, we also add an 8 vs. 8 comparison and for the MontPick data set a 10 vs. 10 comparison, since the larger sample sizes allow this. In this case, we sampled each trial independently so there can be duplicate samples in some cases (that is the comparison of *n*_1_ vs. *n*_2_ might be duplicated. We used this method because it is simple, easy to understand and interpret, and easy to reproduce. If desired, the duplicates could be eliminated with hashing or using the R package partitions (Hankin [63]).

We then compared the number of nominally significant genes with the expected number of significant genes under the null hypothesis. A method with well-calibrated p-values should have a number of significant genes close to the nominal number. With 10,000 genes and 100 replications in each of the two groups, we have 2 million p-values. The expected number less than 10^-4^ (for example) is then 200.

Our methods for comparison differ from previous work in a number of important ways. Soneson and Delorenzi [8], Robles, Qureshi [11], and Guo, Li [13] rely on simulation from parametric distributions (most commonly the negative binomial distribution) for assessing type I error rate and FDR control, which is a useful test only inasmuch as the model assumptions hold on real data. Rapaport, Khanin [14] use technical replicates from the SEQC data set to assess null performance, but do not do a resampling-based comparison. van de Wiel, Neerincx [10] resample from a data set where observations have different covariate values. The approaches used by Reeb and Steibel [12] and Kvam, Liu [9] to evaluate null performance are similar to ours, but Kvam, Liu [9] only report results for FDR-adjusted p-values and Reeb and Steibel [12] do not evaluate performance for unadjusted p-values less than 1e-4. In this paper, we use permutation and resampling as a means of assessing the null performance and power of existing methods. Several methods ([40], [43]) use permutation testing as the basis for analysis. However, Ritchie, Phipson [24] point out, correctly, that permutation testing is hampered by low power under small sample sizes and has limited utility outside of two-group comparisons. We agree with this assessment of permutation testing and believe that methods that control type I error rate without the need for permutation testing pose the only viable alternative.

## 3. Results When There are No True Group Differences

Figure 6 shows a typical result—this is for the MontPick data set with groups of size 3. The pattern is clear: edgeR, edgeR Robust, and DESeq2 have large numbers of false positives, with the disparity growing the smaller the p-value. DESeq has the same pattern, but the curve is translated down, probably by the somewhat artificial use of the maximum of two variance estimates, and there are still large numbers of false positives at the lower p-value thresholds. Limma-voom is very close to on target at all p-value levels. The remainder of the plots for all three data sets and the full range of values of n are to be found in the supplementary material. Limma-voom has some excess false positives for the Bottomly data set at n = 3 and 5 (but not n = 4), though many fewer than the other methods. The results for the Cheung data set are very similar to those of the MontPick data set.

**Figure 6.**
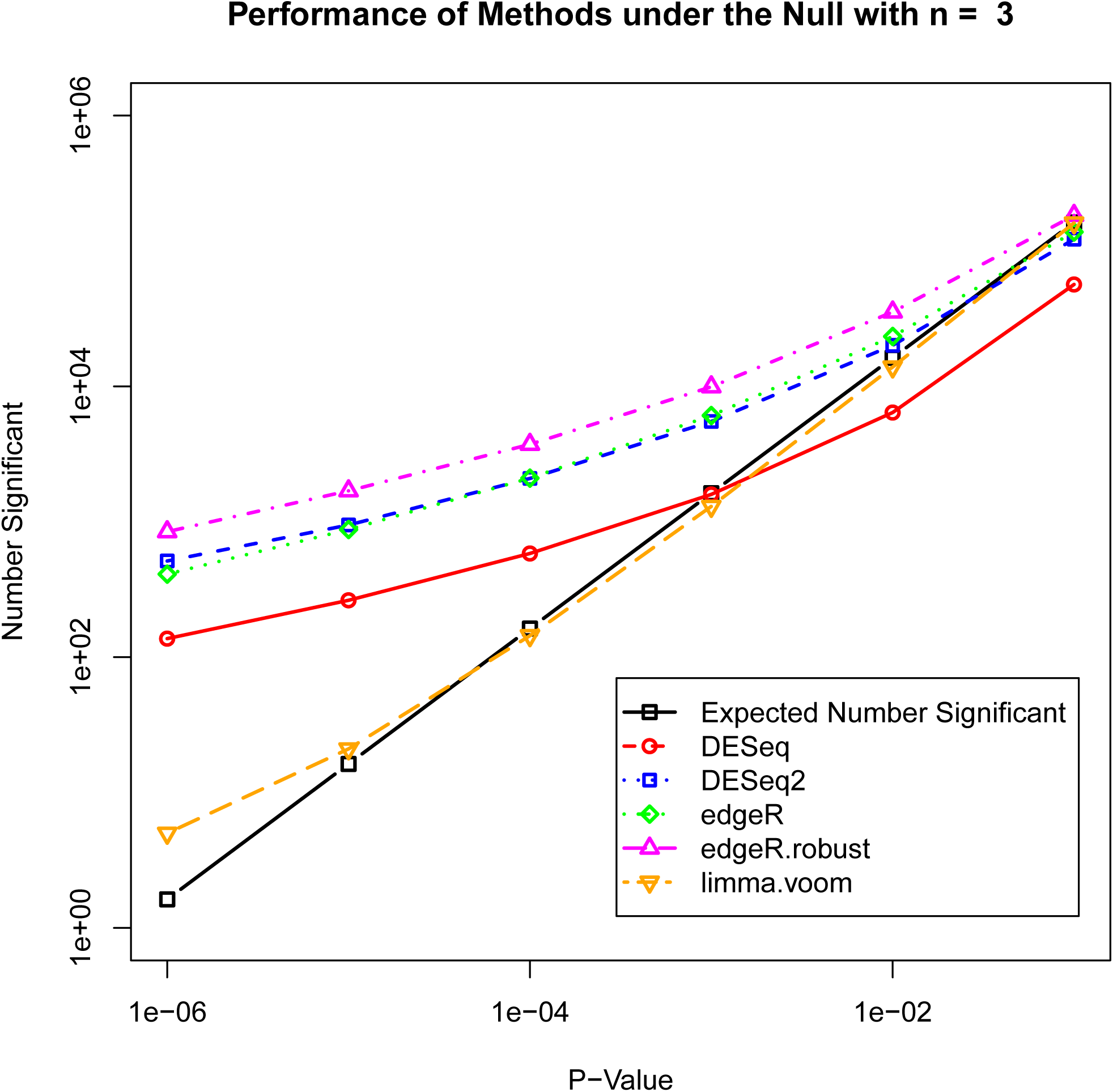
False positives and expected false positives in a null simulation with the MontPick data and n = 3 in each group using DESeq, DESeq2, edgeR, edgeR robust, and limma-voom.

Tables 2 and 3 show the null performance of the five methods on the three data sets at p-value cutoffs of 10^-4^ and 10^-6^. The last five columns show the ratio of the number significant to the expected number significant.

**Table 2.**
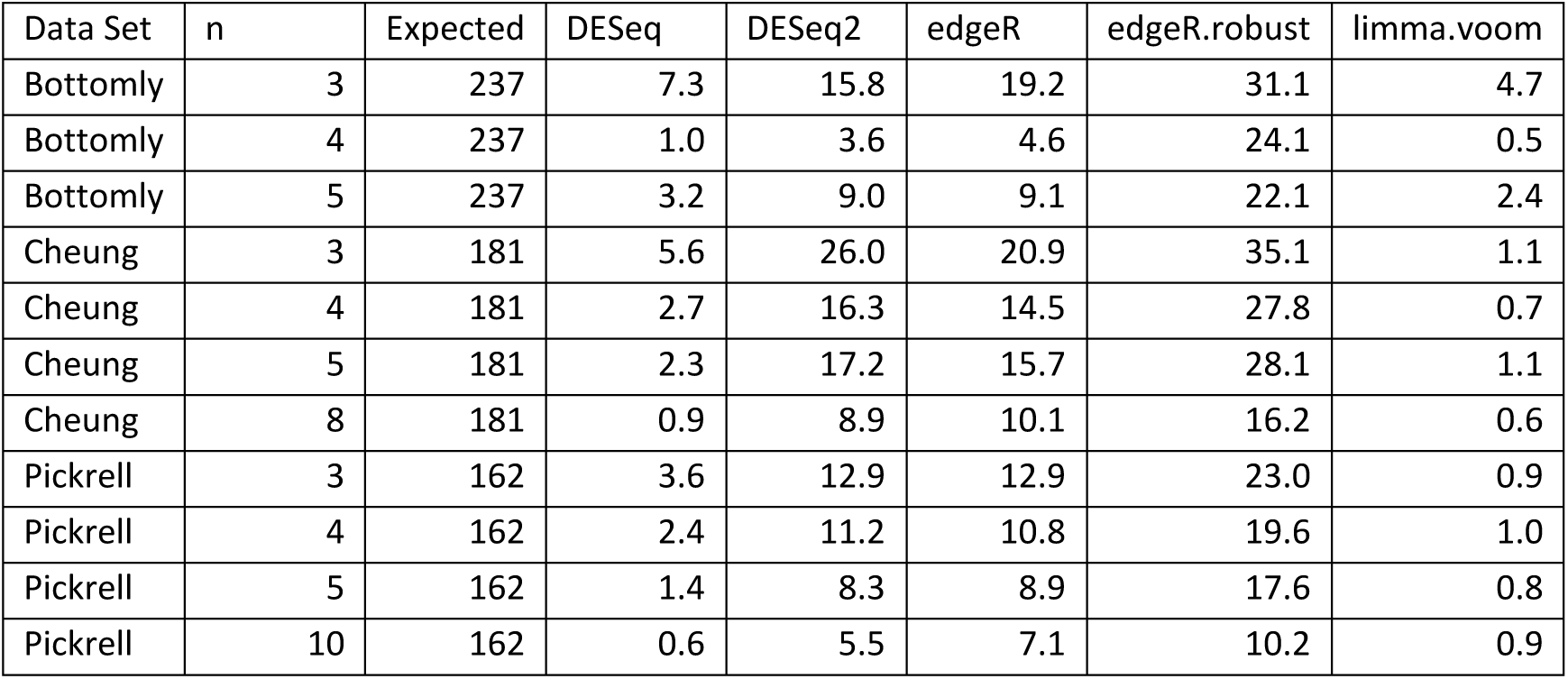
Ratio of number of genes declared significant to number expected for null simulation at a p-value of 10^-4^.

| Data Set | n | Expected | DESeq | DESeq2 | edgeR | edgeR.robust | limma.voom |
| --- | --- | --- | --- | --- | --- | --- | --- |
| Bottomly | 3 | 237 | 7.3 | 15.8 | 19.2 | 31.1 | 4.7 |
| Bottomly | 4 | 237 | 1.0 | 3.6 | 4.6 | 24.1 | 0.5 |
| Bottomly | 5 | 237 | 3.2 | 9.0 | 9.1 | 22.1 | 2.4 |
| Cheung | 3 | 181 | 5.6 | 26.0 | 20.9 | 35.1 | 1.1 |
| Cheung | 4 | 181 | 2.7 | 16.3 | 14.5 | 27.8 | 0.7 |
| Cheung | 5 | 181 | 2.3 | 17.2 | 15.7 | 28.1 | 1.1 |
| Cheung | 8 | 181 | 0.9 | 8.9 | 10.1 | 16.2 | 0.6 |
| Pickrell | 3 | 162 | 3.6 | 12.9 | 12.9 | 23.0 | 0.9 |
| Pickrell | 4 | 162 | 2.4 | 11.2 | 10.8 | 19.6 | 1.0 |
| Pickrell | 5 | 162 | 1.4 | 8.3 | 8.9 | 17.6 | 0.8 |
| Pickrell | 10 | 162 | 0.6 | 5.5 | 7.1 | 10.2 | 0.9 |

**Table 3.** Ratio of number of genes declared significant to number expected for null simulation at a p-value of 10^-6^.

| Data Set | n | Expected | DESeq | DESeq2 | edgeR | edgeR.robust | limma.voom |
| --- | --- | --- | --- | --- | --- | --- | --- |
| Bottomly | 3 | 2.4 | 153.7 | 398.9 | 451.1 | 859.3 | 17.3 |
| Bottomly | 4 | 2.4 | 11.0 | 44.2 | 67.4 | 561.5 | 0.4 |
| Bottomly | 5 | 2.4 | 49.3 | 178.6 | 147.4 | 458.3 | 7.6 |
| Cheung | 3 | 1.8 | 142.8 | 675.0 | 371.7 | 840.4 | 1.1 |
| Cheung | 4 | 1.8 | 50.2 | 295.0 | 193.6 | 539.3 | 0.0 |
| Cheung | 5 | 1.8 | 29.8 | 282.9 | 194.7 | 478.1 | 0.6 |
| Cheung | 8 | 1.8 | 9.4 | 78.9 | 75.0 | 187.5 | 0.0 |
| Pickrell | 3 | 1.6 | 84.3 | 315.7 | 252.3 | 519.4 | 3.1 |
| Pickrell | 4 | 1.6 | 50.5 | 209.9 | 188.9 | 365.0 | 3.1 |
| Pickrell | 5 | 1.6 | 26.5 | 115.7 | 102.2 | 293.6 | 3.1 |
| Pickrell | 10 | 1.6 | 3.1 | 35.1 | 46.2 | 108.3 | 0.6 |

These tables document that edgeR, edgeR Robust and DESeq2 all have inflated false positive rates. DESeq shows this also at smaller p-values, but because of the downward bias caused by the use of the maximum of two variance estimates this is less apparent at larger sample sizes and at higher p-value levels. Limma-voom performs well in nearly every case.

### 3.1 What is the cause of excess false positives?

The question of the cause of the excess false positives could be investigated in a number of ways. The first avenue we explore is whether the fundamental negative binomial assumption is perhaps at the root of the difficulty. We now compare the four standard methods with the worst performance to the use of the likelihood ratio test in the R routine 

~~~
glm.nb()
~~~

, which is a negative binomial generalized linear model routine in the MASS R package. As illustrated in Figure 7, which is a plot of the number of significant genes in the null simulation for the MontPick data with groups of size 3, the likelihood ratio test performs even worse than the other methods. Plots in the supplementary material for the other data sets and sample sizes show that this is no fluke. This means that there is likely to be some aspect of the negative binomial assumption that fails in the actual data and that is at least one cause of the problem is some of the methods.

**Figure 7.**
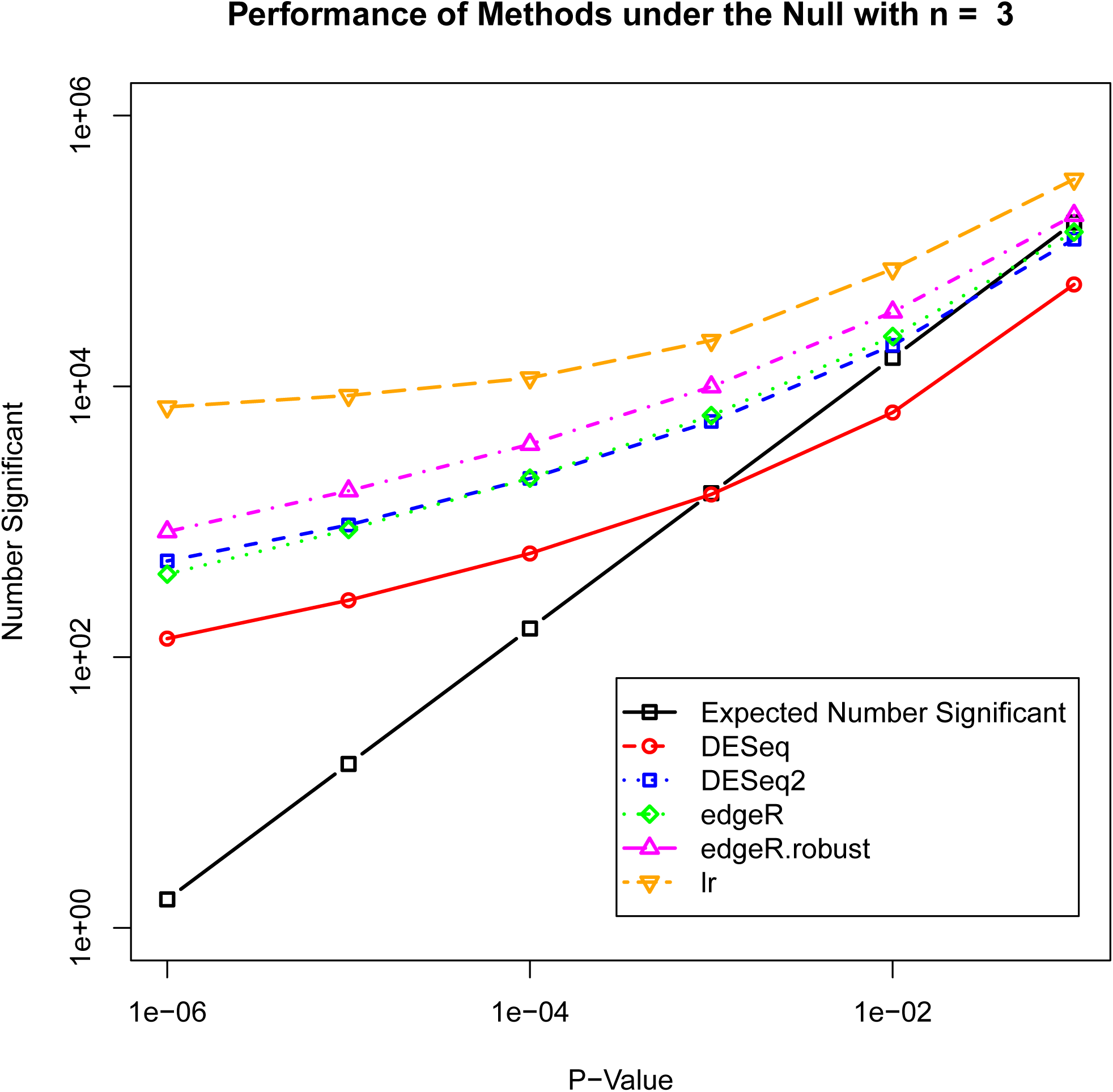
False positives and expected false positives in a null simulation with the MontPick data and n = 3 in each group using DESeq, DESeq2, edgeR, edgeR robust, and the likelihood ratio test in glm.nb().

Another aspect of the problem can be investigated by using additional methods not tied as explicitly to the negative binomial assumption. The first of these is a simple t-test of difference for each gene. The second, which we label limma-trans in the figure, is to employ a variance stabilizing transformation, median normalization, and then use limma without voom pre-processing. Figure 8 shows the same situation as the previous figure but with these three estimators. The t-test is somewhat conservative, but the other two are very close, and they are even closer in larger sample sizes. The remaining cases are shown in figures in the supplementary material and mostly show a similar pattern. The one exception is in the Bottomly data at sample sizes 3 and 5 (but not 4 for some reason). In those cases, limma-voom performs better than the other package methods, but limma-trans has improved performance.

**Figure 8.**
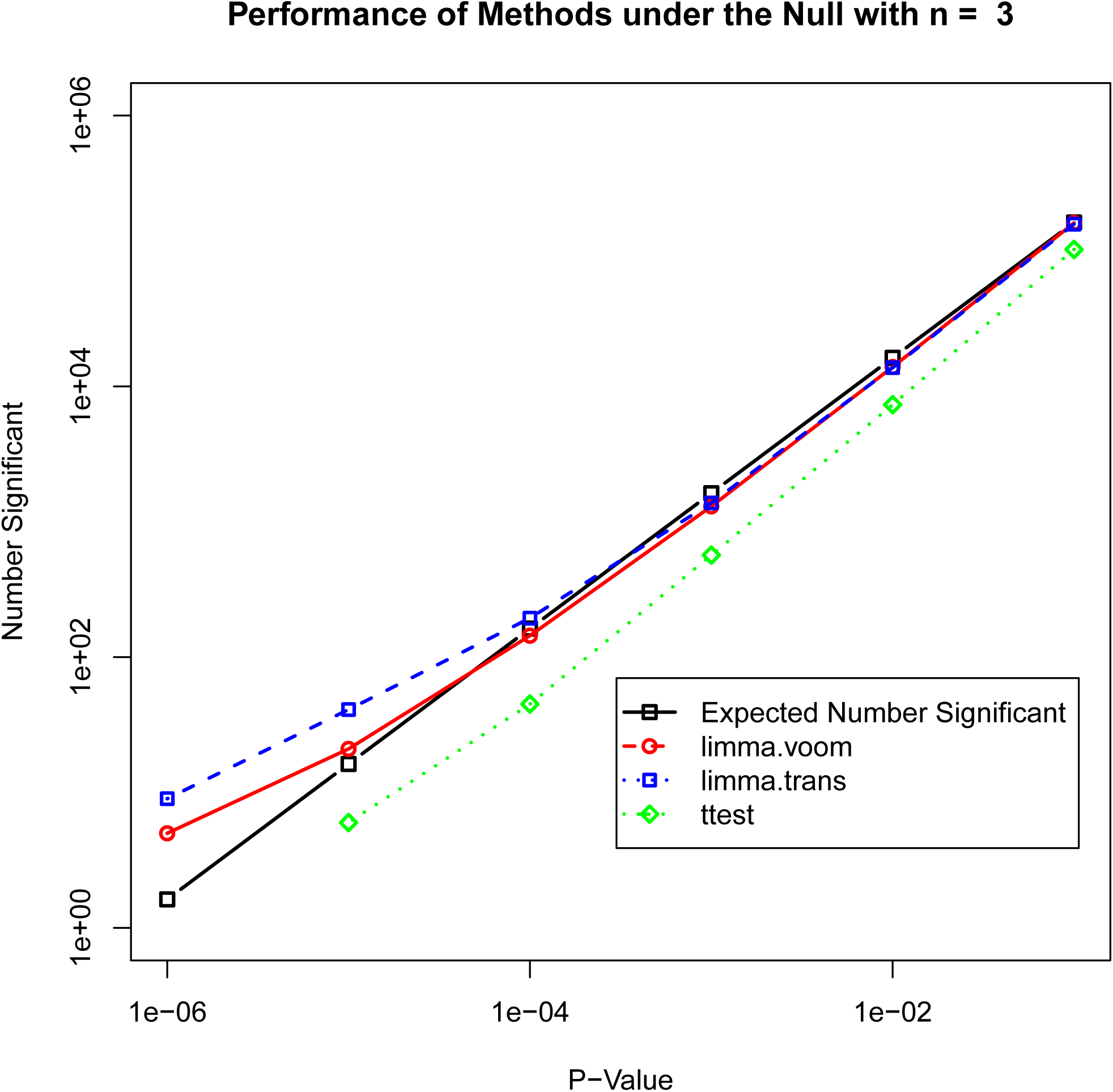
False positives and expected false positives in a null simulation with the MontPick data and n = 3 in each group using limma-voom, limma-trans, and the t-test.

Since the limma documentation is very clear about what it does, we can specify the exact steps performed in the limma-trans method, steps that are not dependent on the limma package, though conveniently performed with it.

1. Transform the data with a variance stabilizing transformation.
2. Normalize (we used median normalization).
3. Estimate a linear model for each gene.
4. Improve the mean square errors using an empirical Bayes method (shrink them towards a middle estimate).
5. Perform statistical tests using the improved mean square error.

Note that the aggregate mean-variance relationship of the negative binomial is used to determine the transformation, but no other negative binomial assumption is used. It should be noted that a method similar to this performed well in the study by Soneson and Delorenzi [8].

## 4. Power

Power in traditional statistical terminology is the probability of rejecting the null hypothesis when it is in fact false. In general, this requires specifying the difference that is intended to be detected. It is also not a meaningful concept if the size of the test is not preserved. So if we want to conduct a hypothesis test with a cutoff for the p-value of 10^-4^, then we cannot meaningfully calculate the power of the test unless the probability of rejection under the null is at least near 10^-4^. So with the methods we have been examining, only limma-voom, limma-trans, and the t-test can be examined for power. Table 4 shows the rejections for analysis of the full MontPick data set in a comparison of the 60 CEU samples with the 69 YRI samples. At every p-value cutoff, the t-test is the least powerful and limma-trans is the most powerful.

**Table 4.** Number of genes significant in a comparison of the 60 CEU samples with the 69 YRI samples in the MontPick data set.

| Significant Genes | limma-voom | limma-trans | t-test |
| --- | --- | --- | --- |
| $p = 10^{-1}$ | 5446 | 5761 | 5280 |
| $p = 10^{-2}$ | 4194 | 4479 | 3875 |
| $p = 10^{-3}$ | 3256 | 3598 | 2900 |
| $p = 10^{-4}$ | 2580 | 2943 | 2238 |
| $p = 10^{-5}$ | 2082 | 2463 | 1742 |
| $p = 10^{-6}$ | 1680 | 2071 | 1377 |

Another way to explore this issue is with non-null simulations. If we select 3 samples from the 60 CEU samples and 3 from the 69 YRI samples, then we can see how many significant genes are obtained at that sample size. Since in the previous section we saw that none of the three methods has excess false positives, this is a measure of power at that sample size. In addition to the previous sample sizes, we looked at n = 20 and n = 30. Figures 9 and 10 show the null simulations for the MontPick data at those samples sizes, showing that excess false positives do not occur. Figures 11–16 show power values for the three methods at the six sample sizes. In every case, limma-trans is somewhat more powerful than limma-voom, and the t-test is the least powerful. This is especially true at small sample sizes because the t-test method we show does not use variance shrinkage.

**Figure 9.**
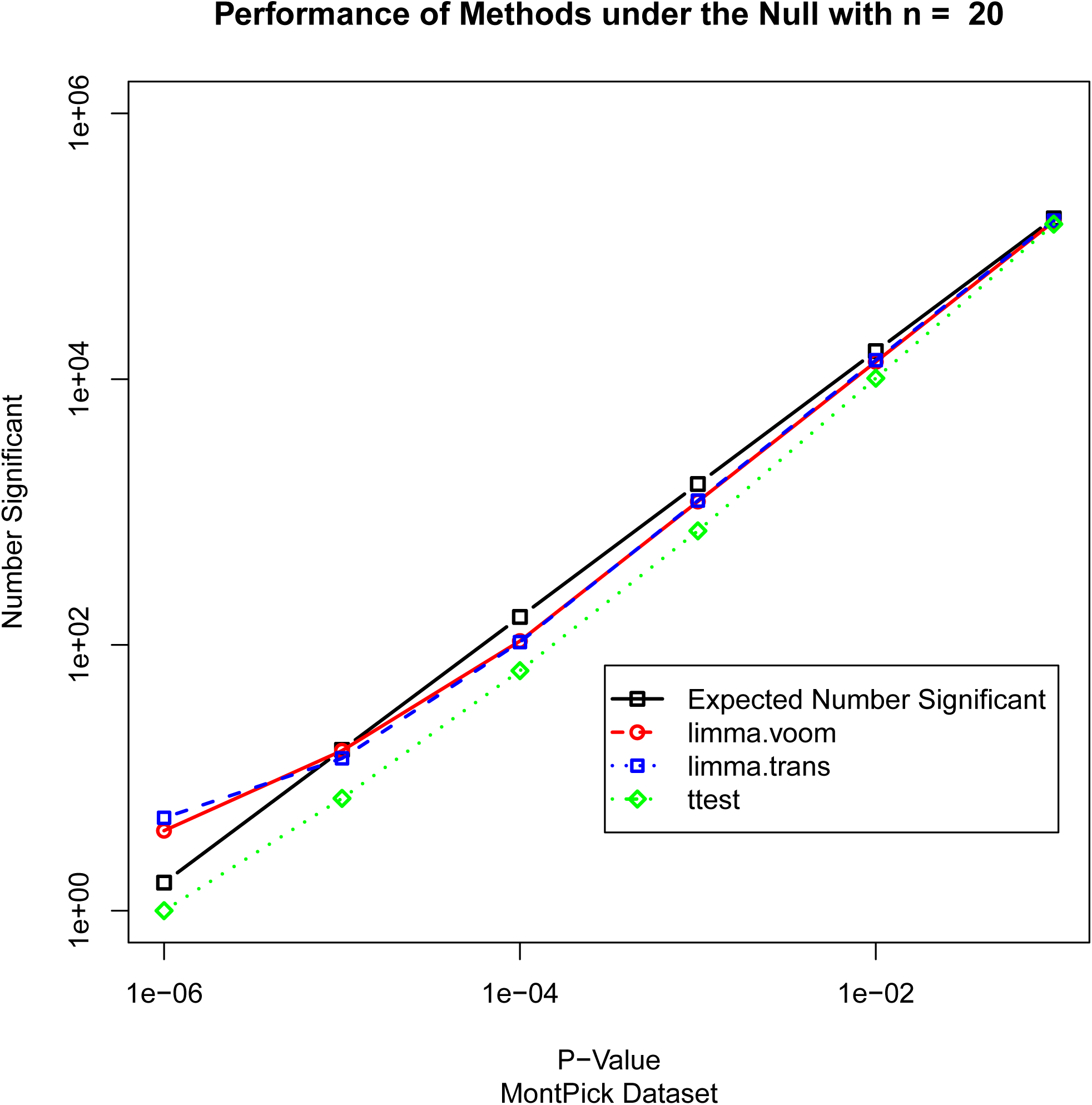
False positives and expected false positives in a null simulation with the MontPick data and n = 20 in each group using limma-voom, limma-trans, and the t-test.

**Figure 10.**
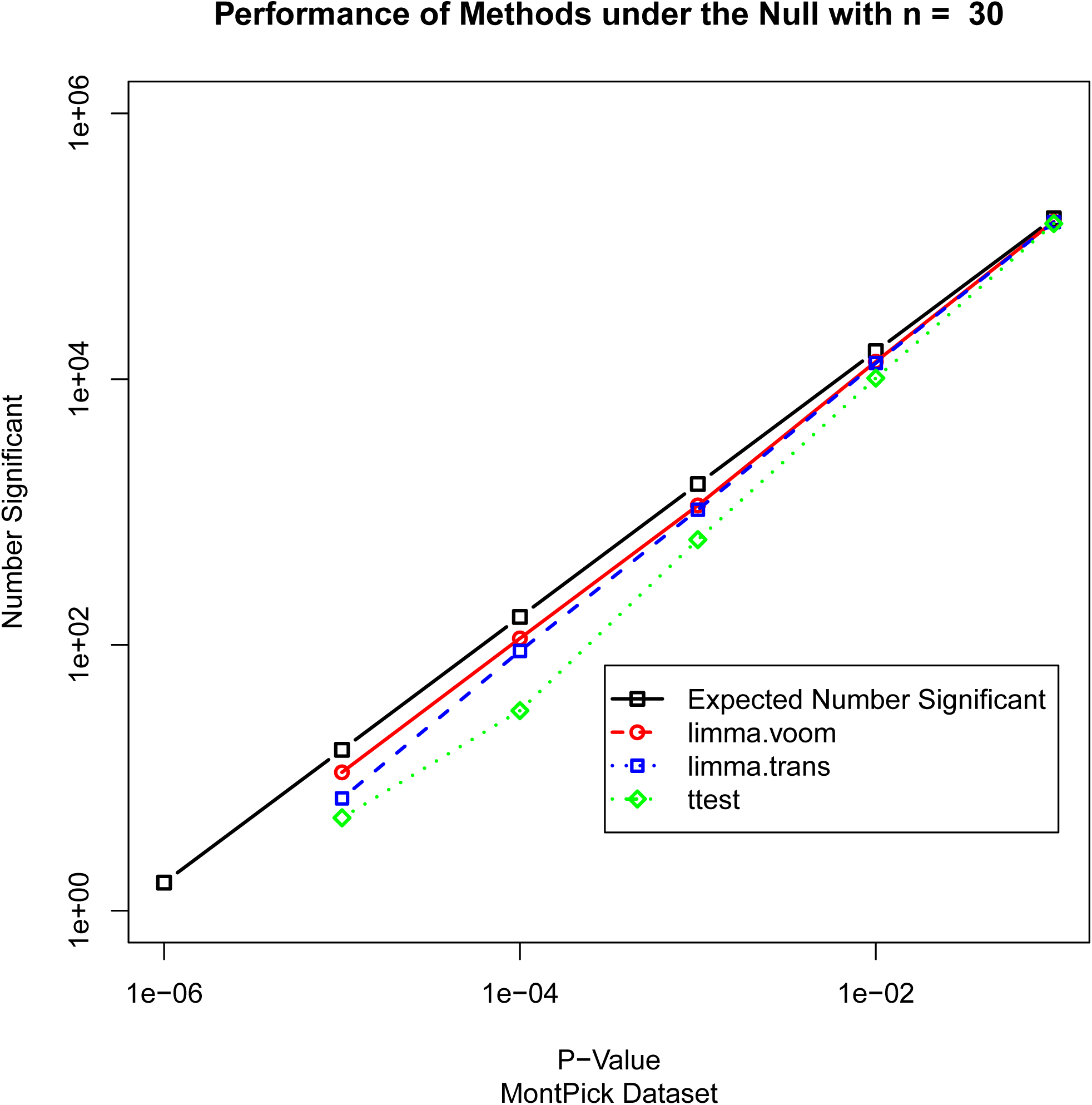
False positives and expected false positives in a null simulation with the MontPick data and n = 30 in each group using limma-voom, limma-trans, and the t-test.

**Figure 11.**
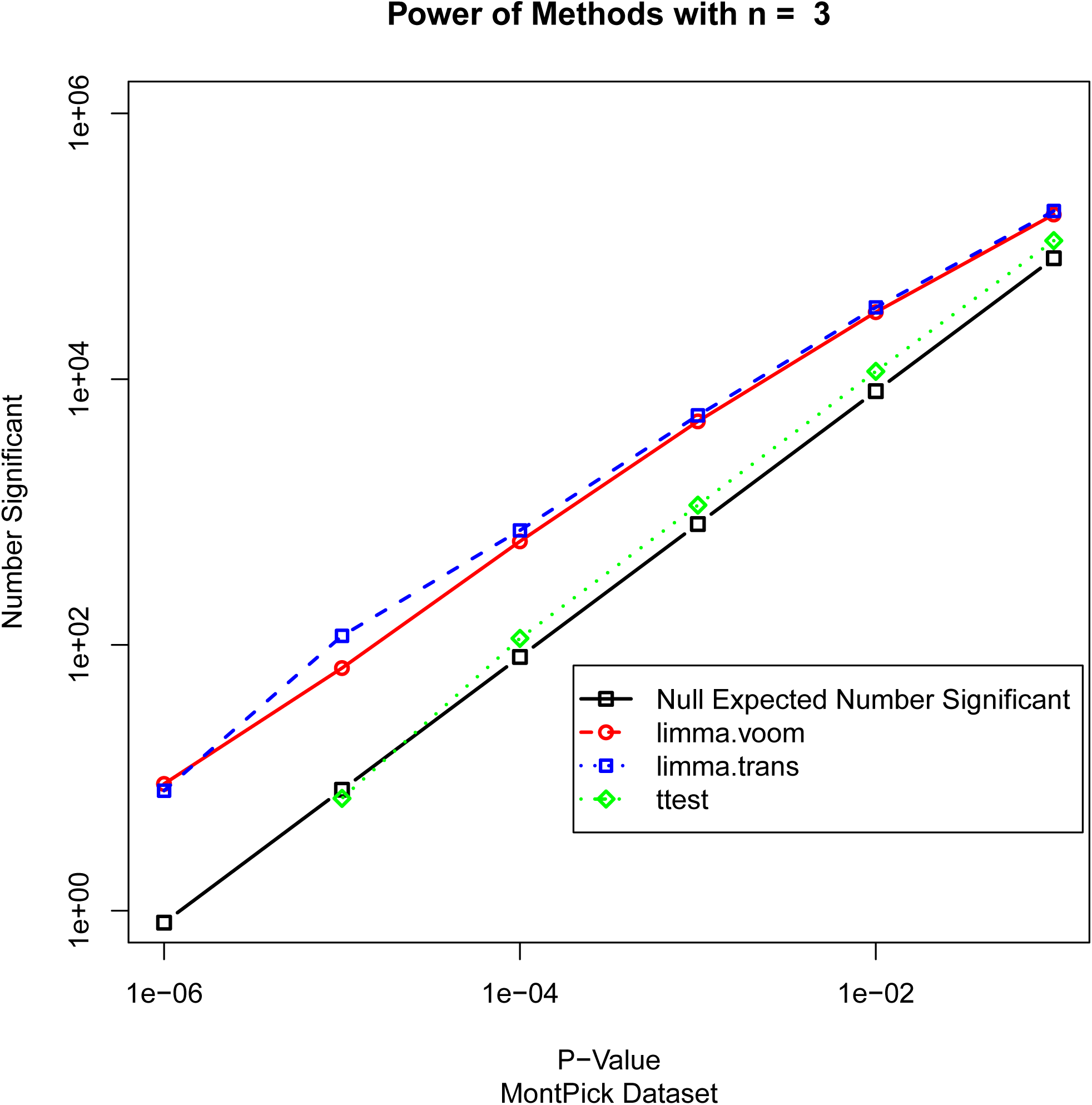
Significant genes in a non-null simulation with the MontPick data and n = 3 in each group using limma-voom, limma-trans, and the t-test.

**Figure 12.**
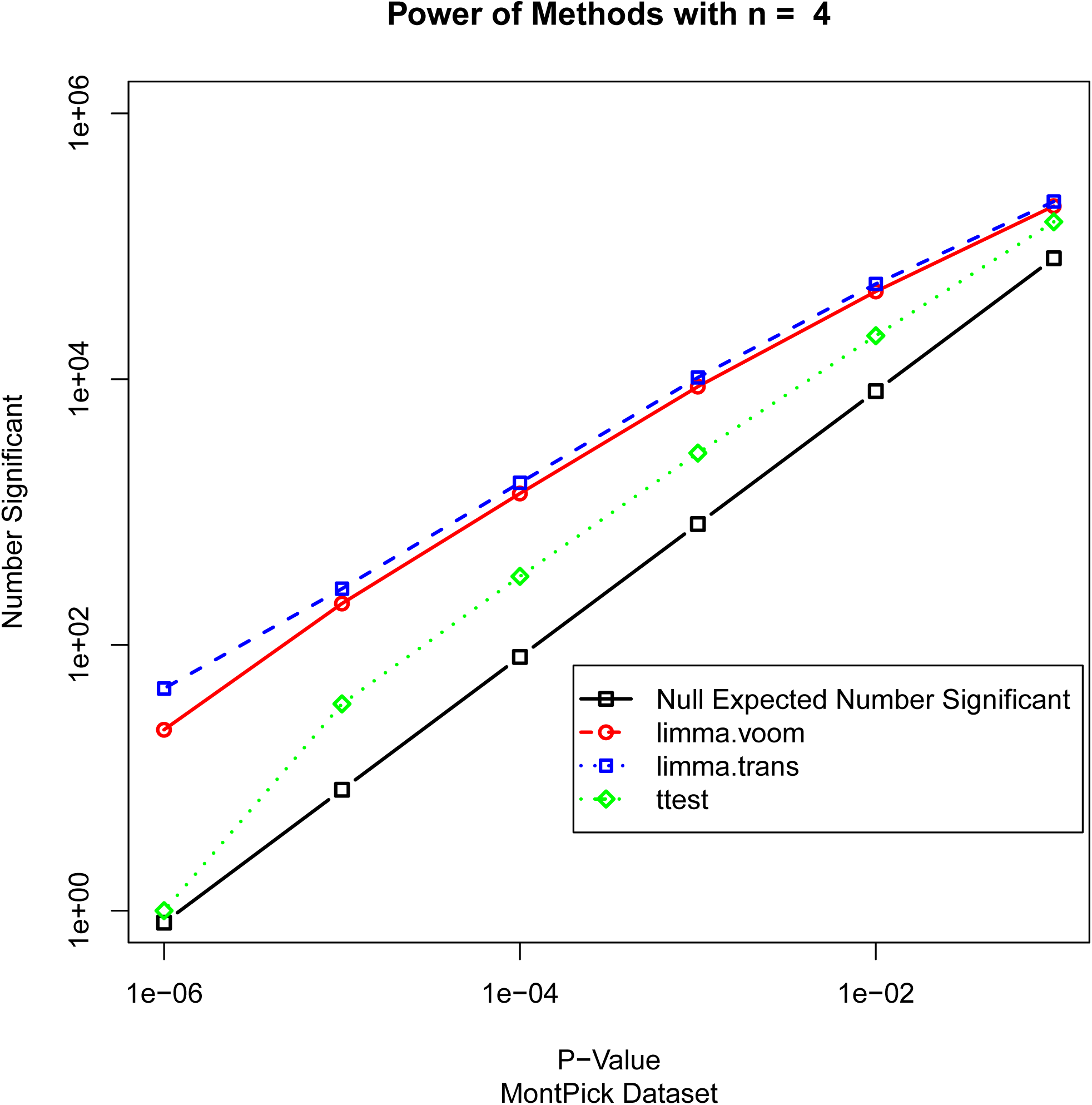
Significant genes in a non-null simulation with the MontPick data and n = 4 in each group using limma-voom, limma-trans, and the t-test.

**Figure 13.**
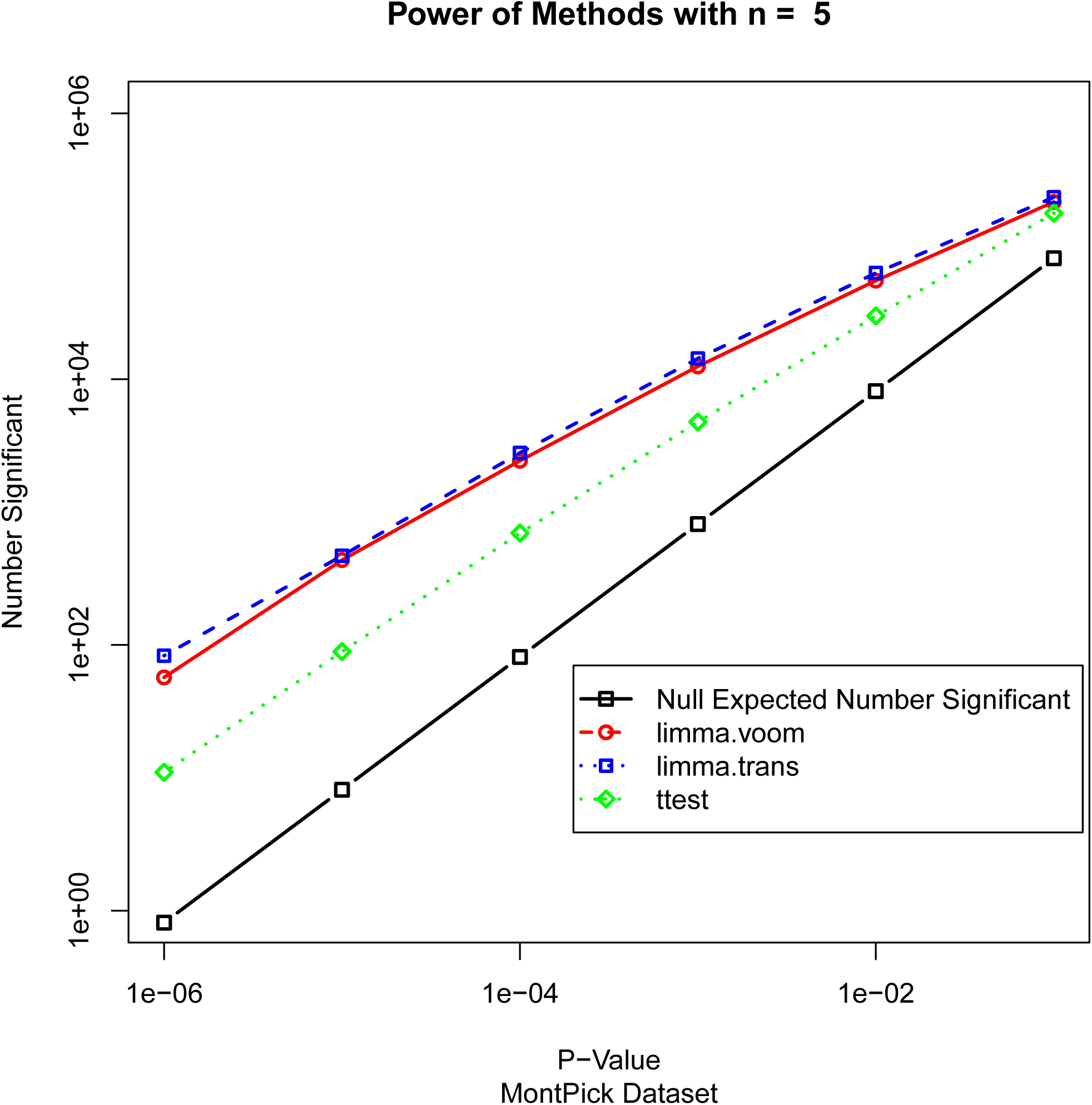
Significant genes in a non-null simulation with the MontPick data and n = 5 in each group using limma-voom, limma-trans, and the t-test.

**Figure 14.**
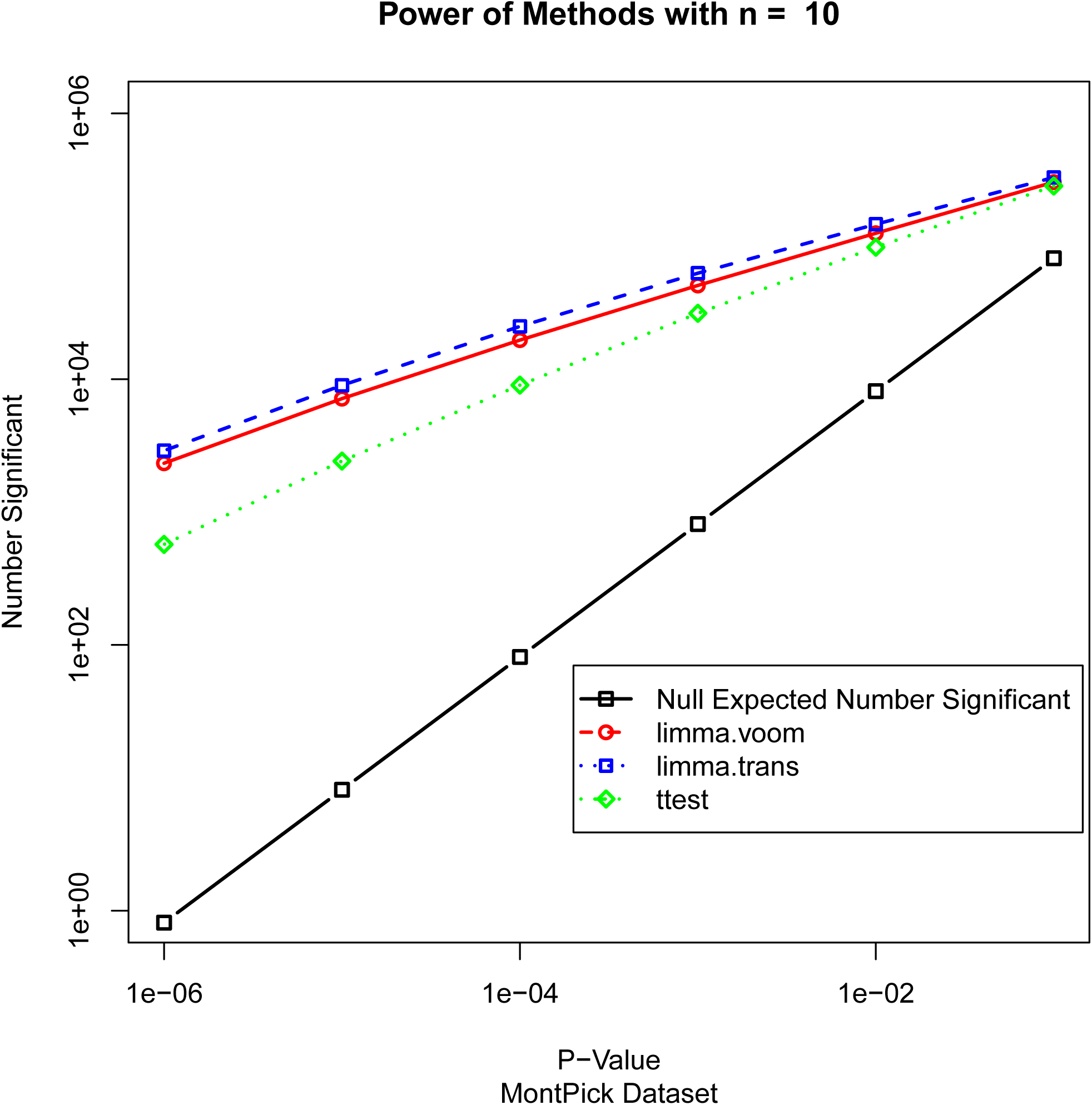
Significant genes in a non-null simulation with the MontPick data and n = 10 in each group using limma-voom, limma-trans, and the t-test.

**Figure 15.**
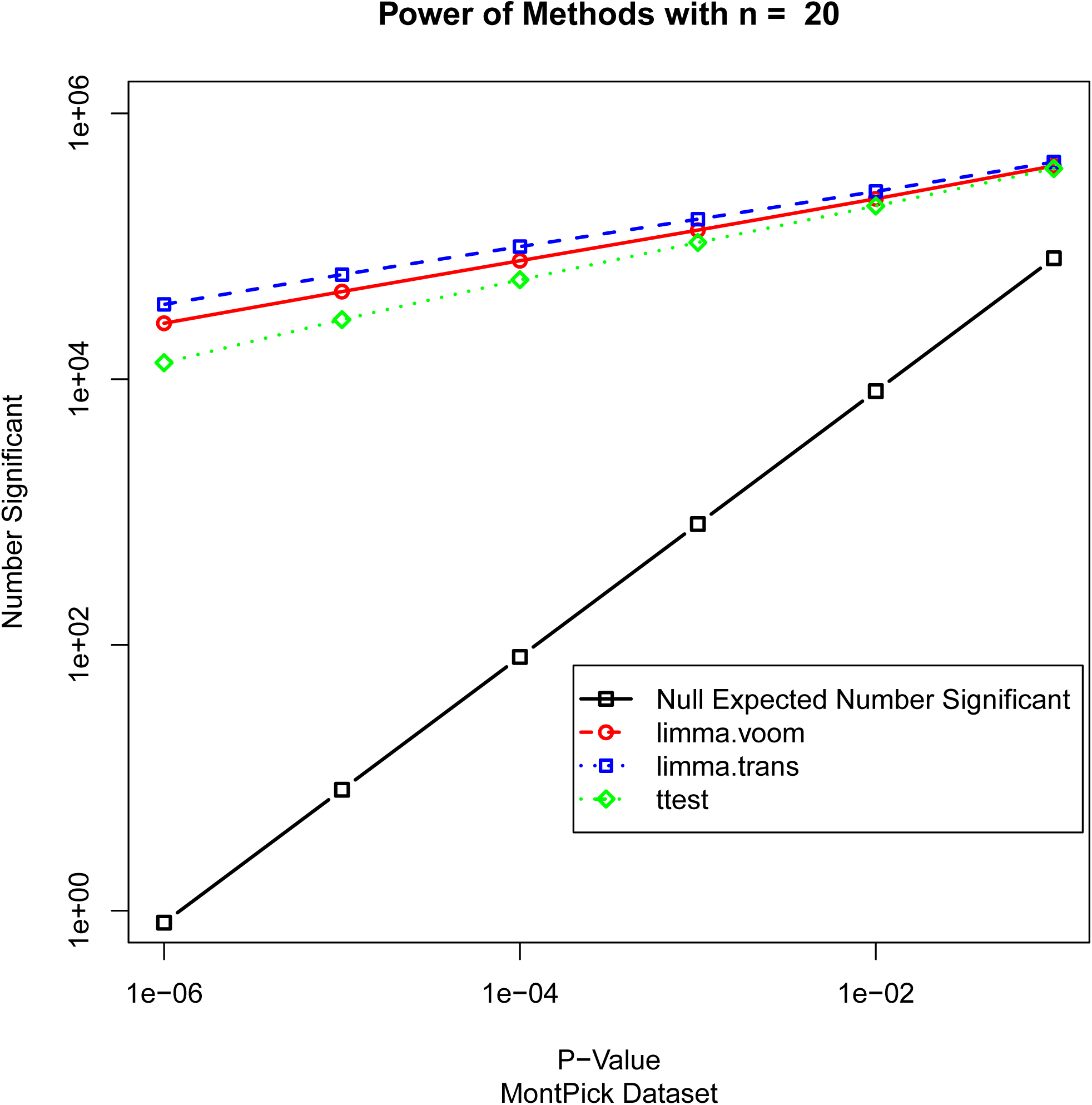
Significant genes in a non-null simulation with the MontPick data and n = 20 in each group using limma-voom, limma-trans, and the t-test.

**Figure 16.**
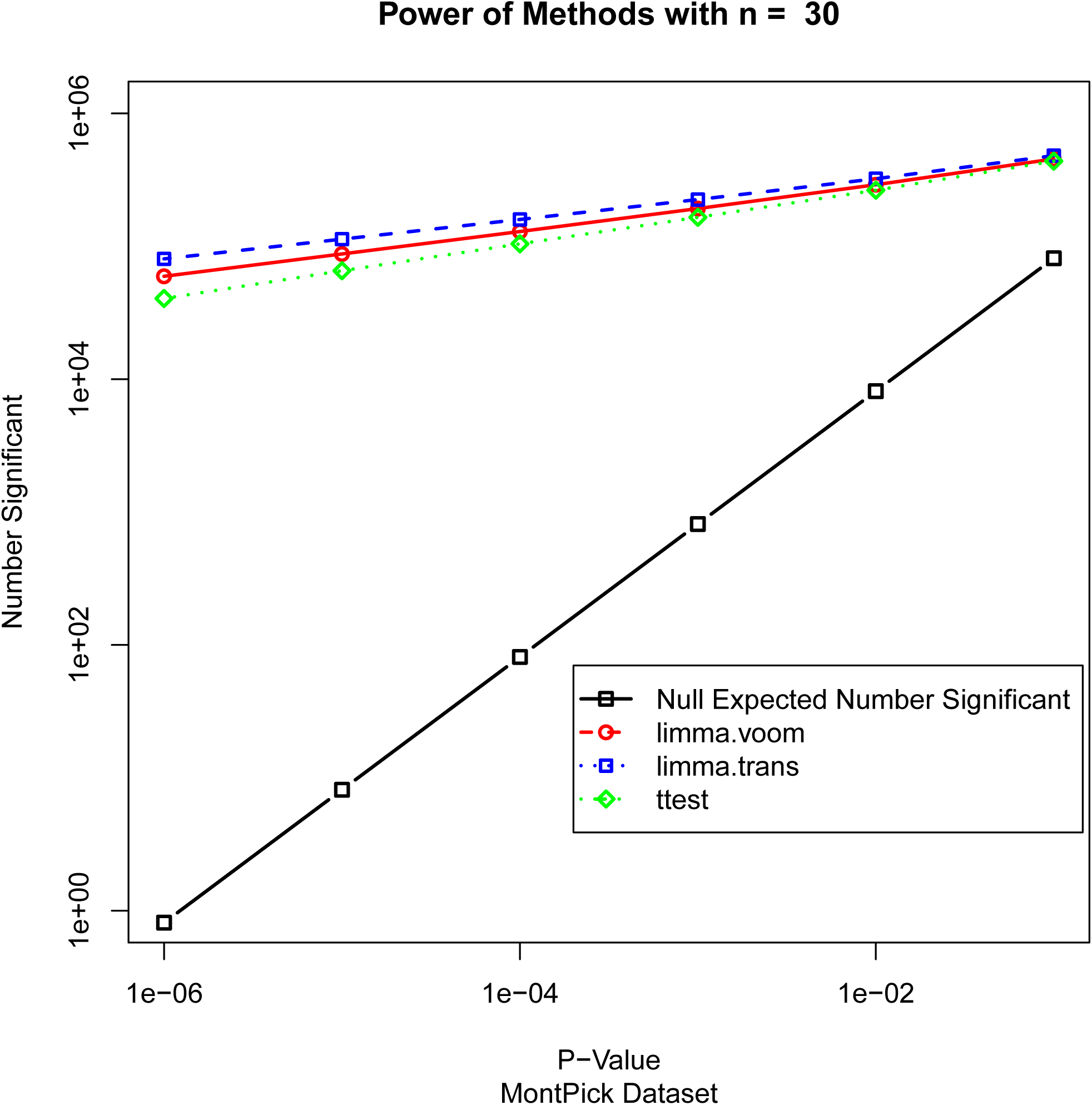
Significant genes in a non-null simulation with the MontPick data and n = 30 in each group using limma-voom, limma-trans, and the t-test.

## 5. Discussion and Conclusions

We have shown that many commonly used packages for differential expression analysis with RNA-Seq data suffer from inflated false-positive rates, while limma-voom and limma with a prior simple data transformation perform well under the null. For the moment, use of limma in one of these two ways seems the safest approach, though the problems with edgeR, DESeq, and DESeq2 may be solvable or improved by tuning the software or by even use of options currently available but not currently the default.

There are several larger points brought up by the analysis in this paper. First, methods matter. Frequently, different methods for the same problem yield similar results, but sometimes for apparent or hidden reasons there is a large performance difference. It is important to know when this happens. It is also often critical to investigate which aspects of a methodology are important and which have little effect on the outcome. For example, voom deals with the large increase of the variance with the mean by weighting the observations, but the transformation method appears to be at least equally effective.

Second, many times methods may become excessively complex, which might obscure the essential nature of the underlying problem and cause unanticipated problems. Certainly the literature as a whole is complex, as evidenced by the 22 packages for RNA-Seq analysis cited in this paper. Whenever possible, we should use the simplest method that works, not the most complex method available. Occam’s razor applies here as well as to scientific hypotheses. Some of the methods we have investigated are straightforward, but others may contain “bells and whistles” that are not necessary.

Third, it may not be necessary to re-invent statistical methodology every time a new type of data arise. The four-step process defined above: transformation, normalization, (possibly) shrinking the variances, and individual statistical tests works equally well for gene expression arrays, proteomics data, metabolomics data and data of other types (Durbin, Hardin [46]; Durbin and Rocke [47]; Durbin and Rocke [48]; Geller, Gregg [64]; Purohit and Rocke [65]; Purohit, Rocke [66]; Rocke [67]; Rocke and Durbin [49]; Rocke and Durbin [50]). The data transformation may differ, but the remainder is very similar.

There are important aspects in the cause of the false positives that still remain to be explored, and this will continue to be an important area of study, so that methods superior to the best of the existing methods can be developed.

### Data accessibility statement

The computational tools developed for this study are freely available via our website http://dmrocke.ucdavis.edu/software.html. They can be downloaded as R code or run directly through an interactive web-based shiny application to reproduce the analysis presented here per a user’s choice of dataset and the methods to be evaluated.

#### Authors’ contributions statement

DMR directed the study, did some of the computations, and was primary author of the manuscript. LR wrote much of the code and performed most of the simulation experiments. JJG wrote some of the code, and did bibliographic research. BDJ and SA wrote parts of the manuscript and edited it and did background research. All participants contributed to the research plan and the outline of the manuscript in extensive and frequent meetings and discussions.

## Footnotes

1 The package versions are as follows: DESeq Package Version 1.18.0; DESeq2 package version 1.6.3; edgeR package Version 3.8.5; and limma package version 3.22.4.

## References

1. Bertone, P., et al., Global identification of human transcribed sequences with genome tiling arrays. Science, 2004. 306(5705): p. 2242–2246.

2. Subramanian, A., et al., Gene set enrichment analysis: a knowledge-based approach for interpreting genome-wide expression profiles. Proceedings of the National Academy of Sciences of the United States of America, 2005. 102(43): p. 15545–15550.

3. Wold, B. and R.M. Myers, Sequence census methods for functional genomics. Nature methods, 2008. 5(1): p. 19–21.

4. Mortazavi, A., et al., Mapping and quantifying mammalian transcriptomes by RNA-Seq. Nature methods, 2008. 5(7): p. 621–628.

5. Nagalakshmi, U., et al., The transcriptional landscape of the yeast genome defined by RNA sequencing. Science, 2008. 320(5881): p. 1344–1349.

6. Van Keuren-Jensen, K., J.J. Keats, and D.W. Craig, Bringing RNA-seq closer to the clinic. Nature biotechnology, 2014. 32(9): p. 884–885.

7. Li, S., et al., Multi-platform assessment of transcriptome profiling using RNA-seq in the ABRF next-generation sequencing study. Nature biotechnology, 2014. 32(9): p. 915–925.

8. Soneson, C. and M. Delorenzi, A comparison of methods for differential expression analysis of RNA-seq data. BMC bioinformatics, 2013. 14(1): p. 91.

9. Kvam, V.M., P. Liu, and Y. Si, A comparison of statistical methods for detecting differentially expressed genes from RNA-seq data. American journal of botany, 2012. 99(2): p. 248–256.

10. van de Wiel, M.A., et al., ShrinkBayes: a versatile R-package for analysis of count-based sequencing data in complex study designs. BMC bioinformatics, 2014. 15(1): p. 116.

11. Robles, J.A., et al., Efficient experimental design and analysis strategies for the detection of differential expression using RNA-Sequencing. BMC genomics, 2012. 13(1): p. 484.

12. Reeb, P.D. and J.P. Steibel, Evaluating statistical analysis models for RNA sequencing experiments. Frontiers in genetics, 2013. 4.

13. Guo, Y., et al., Evaluation of read count based RNAseq analysis methods. BMC genomics, 2013. 4(Suppl 8): p. S2.

14. Rapaport, F., et al., Comprehensive evaluation of differential gene expression analysis methods for RNA-seq data. Genome Biol, 2013. 14(9): p. R95.

15. Nookaew, I., et al., A comprehensive comparison of RNA-Seq-based transcriptome analysis from reads to differential gene expression and cross-comparison with microarrays: a case study in Saccharomyces cerevisiae. Nucleic acids research, 2012: p. gks804.

16. Gentleman, R.C., et al., Bioconductor: open software development for computational biology and bioinformatics. Genome biology, 2004. 5(10): p. R80.

17. Robinson, M.D., D.J. McCarthy, and G.K. Smyth, edgeR: a Bioconductor package for differential expression analysis of digital gene expression data. Bioinformatics, 2010. 26(1): p. 139–140.

18. McCarthy, D.J., Y. Chen, and G.K. Smyth, Differential expression analysis of multifactor RNA-Seq experiments with respect to biological variation. Nucleic acids research, 2012: p. gks042.

19. Robinson, M.D. and G.K. Smyth, Moderated statistical tests for assessing differences in tag abundance. Bioinformatics, 2007. 23(21): p. 2881–2887.

20. Robinson, M.D. and G.K. Smyth, Small-sample estimation of negative binomial dispersion, with applications to SAGE data. Biostatistics, 2008. 9(2): p. 321–332.

21. Zhou, X., H. Lindsay, and M.D. Robinson, Robustly detecting differential expression in RNA sequencing data using observation weights. Nucleic acids research, 2014. 42(11): p. e91–e91.

22. Smyth, G.K., *Limma: linear models for microarray data*, in Bioinformatics and computational biology solutions using R and Bioconductor. 2005, Springer. p. 397–420.

23. Law, C.W., et al., Voom: precision weights unlock linear model analysis tools for RNA-seq read counts. Genome Biol, 2014. 15(2): p. R29.

24. Ritchie, M.E., et al., limma powers differential expression analyses for RNA-sequencing and microarray studies. Nucleic acids research, 2015: p. gkv007.

25. Anders, S. and W. Huber, Differential expression of RNA-Seq data at the gene level–the DESeq package. Heidelberg, Germany: European Molecular Biology Laboratory (EMBL), 2012.

26. Love, M.I., W. Huber, and S. Anders, Moderated estimation of fold change and dispersion for RNA-Seq data with DESeq2. Genome biology, 2014. 15(12): p. 550.

27. Trapnell, C., et al., Transcript assembly and quantification by RNA-Seq reveals unannotated transcripts and isoform switching during cell differentiation. Nature biotechnology, 2010. 28(5): p. 511–515.

28. Trapnell, C., et al., Differential analysis of gene regulation at transcript resolution with RNA-seq. Nature biotechnology, 2013. 31(1): p. 46–53.

29. Di, Y., et al., The NBP negative binomial model for assessing differential gene expression from RNA-Seq. Statistical Applications in Genetics and Molecular Biology, 2011. 10(1): p. 1–28.

30. Auer, P.L. and R.W. Doerge, A two-stage Poisson model for testing RNA-seq data. Statistical applications in genetics and molecular biology, 2011. 10(1): p. 1–26.

31. Hardcastle, T.J. and K.A. Kelly, baySeq: empirical Bayesian methods for identifying differential expression in sequence count data. BMC bioinformatics, 2010. 11(1): p. 422.

32. Leng, N., et al., EBSeq: an empirical Bayes hierarchical model for inference in RNA-seq experiments. Bioinformatics, 2013. 29(8): p. 1035–1043.

33. Tarazona, S., et al., Differential expression in RNA-seq: a matter of depth. Genome research, 2011. 21(12): p. 2213–2223.

34. Li, J. and R. Tibshirani, Finding consistent patterns: A nonparametric approach for identifying differential expression in RNA-Seq data. Statistical methods in medical research, 2013. 22(5): p. 519–536.

35. Van De Wiel, M.A., et al., Bayesian analysis of RNA sequencing data by estimating multiple shrinkage priors. Biostatistics, 2012: p. kxs031.

36. Wang, L., et al., DEGseq: an R package for identifying differentially expressed genes from RNA-seq data. Bioinformatics, 2010. 26(1): p. 136–138.

37. Zhou, Y.-H., K. Xia, and F.A. Wright, A powerful and flexible approach to the analysis of RNA sequence count data. Bioinformatics, 2011. 27(19): p. 2672–2678.

38. Singh, D., et al., FDM: a graph-based statistical method to detect differential transcription using RNA-seq data. Bioinformatics, 2011. 27(19): p. 2633–2640.

39. Li, B. and C.N. Dewey, RSEM: accurate transcript quantification from RNA-Seq data with or without a reference genome. BMC bioinformatics, 2011. 12(1): p. 323.

40. Langmead, B., K.D. Hansen, and J.T. Leek, Cloud-scale RNA-sequencing differential expression analysis with Myrna. Genome Biol, 2010. 11(8): p. R83.

41. Moulos, P. and P. Hatzis, Systematic integration of RNA-Seq statistical algorithms for accurate detection of differential gene expression patterns. Nucleic acids research, 2014: p. gku1273.

42. Fernandes, A.D., et al., Unifying the analysis of high-throughput sequencing datasets: characterizing RNA-seq, 16S rRNA gene sequencing and selective growth experiments by compositional data analysis. Microbiome, 2014. 2(1): p. 1–13.

43. Li, J., et al., Normalization, testing, and false discovery rate estimation for RNA-sequencing data. Biostatistics, 2011: p. kxr031.

44. Srivastava, S. and L. Chen, A two-parameter generalized Poisson model to improve the analysis of RNA-seq data. Nucleic acids research, 2010. 38(17): p. e170–e170.

45. Anders, S. and W. Huber, Differential expression analysis for sequence count data. Genome biol, 2010. 11(10): p. R106.

46. Durbin, B.P., et al., A variance-stabilizing transformation for gene-expression microarray data. Bioinformatics, 2002. 18(suppl 1): p. S105–S110.

47. Durbin, B. and D.M. Rocke, Estimation of transformation parameters for microarray data. Bioinformatics, 2003. 19(11): p. 1360–1367.

48. Durbin, B.P. and D.M. Rocke, Variance-stabilizing transformations for two-color microarrays. Bioinformatics, 2004. 20(5): p. 660–667.

49. Rocke, D.M. and B. Durbin, A model for measurement error for gene expression arrays. Journal of computational biology, 2001. 8(6): p. 557–569.

50. Rocke, D.M. and B. Durbin, Approximate variance-stabilizing transformations for gene-expression microarray data. Bioinformatics, 2003. 19(8): p. 966–972.

51. Welch, B.L., The generalization ofstudent’s’ problem when several different population variances are involved. Biometrika, 1947: p. 28–35.

52. Ripley, B., MASS: support functions and datasets for Venables and Ripley’s MASS. R package version, 2011: p. 7.3–29.

53. Neyman, J. and E.S. Pearson, On the use and interpretation of certain test criteria for purposes of statistical inference: Part I. Biometrika, 1928: p. 175–240.

54. Snedecor, G.W., Calculation and interpretation of analysis of variance and covariance. 1934.

55. Bartlett, M.S., The use of transformations. Biometrics, 1947. 3(1): p. 39–52.

56. Pickrell, J.K., et al., Understanding mechanisms underlying human gene expression variation with RNA sequencing. Nature, 2010. 464(7289): p. 768–772.

57. Montgomery, S.B., et al., Transcriptome genetics using second generation sequencing in a Caucasian population. Nature, 2010. 464(7289): p. 773–7.

58. Wright, G.W. and R.M. Simon, A random variance model for detection of differential gene expression in small microarray experiments. Bioinformatics, 2003. 19(18): p. 2448–2455.

59. Smyth, G.K., Linear models and empirical bayes methods for assessing differential expression in microarray experiments. Statistical applications in genetics and molecular biology, 2004. 3(1): p. 1–25.

60. Baldi, P. and A.D. Long, A Bayesian framework for the analysis of microarray expression data: regularized t-test and statistical inferences of gene changes. Bioinformatics, 2001. 17(6): p. 509– 519.

61. Bottomly, D., et al., Evaluating gene expression in C57BL/6J and DBA/2J mouse striatum using RNA-Seq and microarrays. PloS one, 2011. 6(3): p. e17820.

62. Cheung, V.G., et al., Polymorphic cis- and trans-regulation of human gene expression. PLoS Biol, 2010. 8(9).

63. Hankin, R.K., Additive integer partitions in R. Journal of Statistical Software, 2006. 16(Code Snippet 1): p. 3pp.

64. Geller, S.C., et al., Transformation and normalization of oligonucleotide microarray data. Bioinformatics, 2003. 19(14): p. 1817–1823.

65. Purohit, P.V. and D.M. Rocke, Discriminant models for high-throughput proteomics mass spectrometer data. Proteomics, 2003. 3(9): p. 1699–1703.

66. Purohit, P.V., et al., Discrimination models using variance-stabilizing transformation of metabolomic NMR data. Omics: a journal of integrative biology, 2004. 8(2): p. 118–130.

67. Rocke, D.M. Design and analysis of experiments with high throughput biological assay data. in Seminars in cell & developmental biology. 2004. Elsevier.

